# Design, synthesis and biological evaluation of novel quercetin derivatives as PPAR-γ partial agonists by modulating Epithelial-mesenchymal transition in lung cancer metastasis

**DOI:** 10.1101/2022.09.11.507444

**Authors:** Sangeeta Ballav, Mrinalini Bhosale, Kiran Bharat Lokhande, Manash K. Paul, Subhash Padhye, K. Venkateswara Swami, Amit Ranjan, Soumya Basu

## Abstract

Epithelial-to-mesenchymal transition (EMT) is responsible for driving metastasis of multiple cancer types including lung cancer. Peroxisome proliferator-activated receptor (PPAR)-γ, a ligand-activated transcription factor, controls expression of variety of genes involved in EMT, cellular differentiation, fatty acid metabolism, insulin sensitivity and adipogenesis. Several synthetic compounds act as potent full agonist for PPAR-γ. However, owing to their serious adverse effects, restricts their long-term application. Therefore, partial agonist has been greatly in demand which involves reduced and balanced PPAR-γ activity. Our previous study discerned the efficacy of quercetin and its derivatives to attain favourable stabilization with PPAR-γ. Here we extended this work by synthesizing five novel quercetin derivatives (QDs) namely thiosemicarbazone (QUETSC) and hydrazones (QUEINH, QUENH, QUE2FH and QUESH) and analysed their effects in modulating EMT of lung cancer cell lines via PPAR-γ partial activation. QDs-treated A549 cells exhibited cell death strongly in a dose and time dependent manner at nanomolar concentration along with anti-migratory effects compared to NCI-H460 cells. Of the five derivatives we screened, QUETSC, QUE2FH and QUESH exhibited the property of partial activation as compared to the over-expressive level of rosiglitazone (RSG). Consistently, with PPAR-γ partial activation, these QDs also suppressed EMT process by markedly down-regulating the levels of mesenchymal markers (Snail, Slug and Zeb-1) and concomitant up-regulation of epithelial marker (E-cadherin). In the light of these evidences; QUETSC, QUE2FH and QUESH could be used as a novel selective partial PPAR-γ modulators whose pharmacological properties is distinct from RSG and may be exploited as potential therapeutic anti-metastatic agent.

## INTRODUCTION

Lung cancer have been identified as one of the serious causes of death worldwide, accounting 85% of the cases to be diagnosed with non-small-cell lung cancer (NSCLC) (Siegel *et al*. 2022). According to the mortality pattern of National Center for Health Statistics, death rates are more accelerating for lung cancer as compared to other cancers (SEER*Explorer 2021).This underscores to develop innovative therapies for early detection, prevention, and treatment of this disease. Epithelial–mesenchymal transition (EMT) is a highly dynamic process that has a central role in tumor progression. EMT is characterized by transition of polarized epithelial cells to motile mesenchymal cells, leading to manifested biological morphogenetic process (Yang *et al*. 2020).Intense research efforts in the last two decades revealed that EMT mechanism can be modulated upon the activation of Peroxisome Proliferator-activated Receptor-γ (PPAR-γ) which is a prominent tumor-suppressive player (Li *et al*. 2013; Reka *et al*. 2010).

PPAR-γ is a nuclear receptor family protein that processes the transcription of various target genes in EMT on its activation by certain ligands (Choi *et al*. 2014). An early study conducted by Tan *et al*. have illustrated the inhibition of pro-fibrotic changes in A549 cells in TGF-β1-induced EMT process through the activation of PPAR-γ by its potent synthetic ligands (thiazolidinediones (TZDs)), namely, rosiglitazone and ciglitazone (Tan et al. 2010). Although synthetic ligands act as strong activators of PPAR-γ, the application has been restricted due to their toxicity profile viz. fluid retention, weight gain and hepatotoxicity (Nesto *et al*. 2003; Guan *et al*. 2005). Therefore, a moderate activation may serve to be a novel and intriguing therapeutic strategy which would encompass reduced and balanced PPAR-γ activity. Our previous *in silico* findings discerned that quercetin and its derivatives were observed to exert minimal conformational changes and attained a favourable stabilization with PPAR-γ (Lokhande et al. 2020; Lokhande *et al*. 2020). Quercetin, a naturally occurring polyphenolic flavonoid, marks a great virtue of beneficial effects along with low tissue toxicity and anti-cancer properties (Tang *et al*. 2020). In addition, recent findings have also highlighted the significant role of quercetin in restraining EMT (Chuang *et al*. 2016; Lu *et al*. 2020; Ravichandran *et al*. 2014). In an instance, Kim SR and colleagues explored cytotoxic and metastatic effects of oral squamous cancer cells by reversing the action of EMT induced by TGF-β1 where they observed the attenuation of mesenchymal markers and decreased MMP-2 and MMP-9 expression on administration of 40 µM (IC_50_) of quercetin (Kim *et al*. 2020).

More likely, quercetin has been reported to act as partial weak agonist for PPAR-γ.Fang *et al*. proved the efficiency of quercetin metabolites in stimulating PPAR-γ expression and in turn inhibiting A549 cell growth with downregulation of cdk1 and cyclin B expression (Fang *et al*.2008).Therefore, we sought to build novel and better partial agonist taking quercetin as template to improvise the overall therapeutics as implicated from our earlier data (Lokhande et al. 2020; Lokhande *et al*. 2020).Thusly, to accomplish our objective, in the present work, we have synthesized quercetin derivatives with the introduction of Schiff base compounds, namely, Thiosemicarbazone (TSC) (RNH–CS–NH–N=CR^1^R^2^) and Hydrazone (R^1^R^2^C=N-NH^2^). They have immense biological importance, high synthetic flexibility and increased resemblance for natural compounds which offers promiscuous notion for constructing novel biological active structures (Afzal *et al*. 2021; Chen *et al*. 2011).The synthesized compounds were characterized via chromatographic and spectrometric techniques along with predicting their absorption, distribution, metabolism, and excretion (ADME) properties. They were then analysed for their interaction with PPAR-γ to unravel the underlying the binding affinity through the process of molecular docking and molecular dynamic (MD) simulation. Furthermore, the effects of these synthesized quercetin derivatives (QDs) were investigated on two NSCLC cell lines-A549 and NCI-H460 for their anti-proliferative and anti-migratory abilities. In addition, quercetin also ought to act hold upon migration and metastasis by modulating EMT signalling components (Fang *et al*.2008; Lu *et al*. 2020). Although quercetin inhibits tumor progression in lung cancer, the detailed mechanisms are not fully understood. Thereafter, the QDs were examined for their partial agonist activity towards PPAR-γ pathway through western blotting and transcription factor binding assay which were in turn analyzed for understanding their effect on EMT modulation. The intent of the study is to yield better potential PPAR-γ partial agonist which might substantially modulate EMT of NSCLC cell lines via the partial activation. Thus, this would throw light on reviving the therapeutic condition of lung cancer treatment via PPAR-γ.

## EXPERIMENTAL

### Materials

Quercetin and Rosiglitazone (RSG) were purchased from Cayman Chemical Company (MI, USA). For chemical synthesis, all solvents and reagents were procured from commercial sources; Sigma Aldrich (MO, USA), Thermo Fischer Scientific (MA, USA), Santa Cruz Biotechnology (TX, USA), and Merck (NJ, USA), which were used without further purification. Stock solutions of QDs and quercetin were prepared in dimethyl sulfoxide (DMSO), whereas RSG was dissolved in water and used at a final concentration of 1mM. Roswell Park Memorial Institute Medium (RPMI)-1640 medium, 200 U/mL of penicillin, 0.2 mg/ml of streptomycin and 200mM L-glutamine, 10% fetal bovine serum (FBS), 1% glutamine (200 mM), 0.25% trypsin/0.038% EDTA, crystal violet stain, 3-(4,5-dimethythiazol2-yl)-2,5-diphenyl tetrazolium bromide (MTT) dye was purchased from HiMedia (Mumbai, India). TRIzol reagent was purchased from Ambion, life technologies. Enhanced chemiluminescent (ECL) substrate kit was purchased from Bio-rad (CA, USA).Antibodies details are as follows; Mouse monoclonal anti-human β-actin antibody and rabbit polyclonal anti-human PPAR-γ antibody was obtained from Cloud clone, EMT markers specific antibodies such as-rabbit anti-Snail, anti-N-cadherin, anti-Vimentin, anti-Slug, anti-E-cadherin and anti-Zeb-1 antibodies were obtained from Cell Signaling Technology, CA, USA. Anti-rabbit and anti-mouse antibodies conjugated to horseradish peroxidase (HRP) were also obtained from Cell Signaling Technology, CA, USA.

### General chemistry methods

Melting points (M.P.) were determined using a traditional set up and expressed in degree Celsius (°C). The Fourier Transform Infrared Spectrometer (FT-IR) spectroscopy for all the compounds were run in JASCO FT/IR-4200 spectrophotometer with potassium bromide (KBr) pellets (9:1) and expressed in cm^-1^. NMR spectra were recorded using BrukerAVANCE III HD NMR 500MHz operating at a magnet capacity of 11.7 Tesla. The chemical shifts are provided in parts per million (ppm) with reference to the respective solvent peak, and coupling constants (J) are reported in Hz. The spin multiplicities are given as s-singlet, d-doublet, dd-double doublet, t-triplet, m-multiplet, and br-broad. The reaction progress was monitored using thin-layer chromatography (TLC) (chloroform:methanol, 9:1) on 0.25 mm silica gel plates (60F 254). The purity (> 95%) of the derivatives was monitored by High-performance liquid chromatography (HPLC) experiment using Systronics HPLC (SYS LC-138).

### General procedure for synthesis of QDs

The chemical synthesis of the derivatives was carried out at Prof. Subhash Padhye’s Laboratory, Interdisciplinary Sciences and Technology Research Area Academy (ISTRA), University of Pune, India. Each derivatives (thiosemicarbazine (100mg); Isonicotinic acid hydrazine (QUEINH) (130mg); Nicotinic acid hydrazine (QUENH) (130mg); 2-Furoic hydrazine (QUE2FH) (120mg) and Salicyl hydrazine (QUESH) (120mg)) were mixed with 300mg of Quercetin separately and dissolved in 20mL ethanol as a solvent. 50 µL sulfuric acid (H_2_SO_4_) was added, which served as a catalyst. Each of these reactions were refluxed by heating and continuous stirring for 24 h at 65°C, followed by filtering of the products through a Whatmann filter paper (no. 41) until TLC showed no product in the aqueous phase. Upon completion of the filtration, the residual mass was allowed to dry and evaporate under reduced pressure to get a crude product that was further purified by performing HPLC.

### QUETSC

Colour-Yellow; M.P.-370°C; Mol wt.-375.36g/mol; t_R_-5.615 min;FT-IR (cm^−1^):1744.00 (C=O), 1319.00 (Aromatic C–H), 878.00, 707.27, 637.00 (Aromatic C-H bending), 934.00 (N-N), 3290.00 (OH stretching), 1614.00 (Aromatic C=C stretching for B-ring and C-ring), 1360.00 (OH bending of phenol); ^1^H NMR (DMSO, 500 MHz, ppm)-5.8(d),6.69(s), 6.51(s), 6.60(s), - OH(5.0),-NH(7.0),-NH_2_(2.0).

### QUEINH

Colour-Light Yellow; M.P.-367°C; Mol wt.-421.38 g/mol; t_R_-5.617 min; FT-IR (cm^−1^): 1701.00 (C=O), 1247.89 (Aromatic C–H), 858.00, 755.43, 665.63 (Aromatic C-H bending), 952.64 (N-N), 3093.00 (OH stretching), 1587.00 (Aromatic C=C stretching for B-ring and C-ring), 1330.00 (OH bending of phenol);^1^H NMR(DMSO, 500 MHz, ppm)-5.8(d),6.69(s), 6.51(s), 6.60(s), 9.06(d), 7.96(d), -OH(5.0),-NH(7.0), OH(15.0).

### QUENH

Colour-Light Yellow; M.P.-372°C; Mol wt.-421.36 g/mol; t_R_-5.613 min; FT-IR (cm^−1^): 1710.00 (C=O), 1260.43 (Aromatic C–H), 813.11, 795.00, 694.13 (Aromatic C-H bending), 980.00 (N-N), 3163.13 (OH stretching), 1512.00 (Aromatic C=C stretching for B-ring and C-ring), 1365.11 (OH bending of phenol);^1^H NMR (DMSO, 500 MHz, ppm)-5.8(d),6.69(s), 6.51(s), 6.60(s),8.32(s), 7.63(s), 8.84(s), 9.17(s),-OH(5.0),-OH(15.0).

### QUE2FH

Colour-Orange; M.P.-373°C; Mol wt.-410.33 g/mol; t_R_-5.615 min; FT-IR (cm^−1^): 1655.00 (C=O), 1312.99 (Aromatic C–H), 760.40, 703.45, 574.00 (Aromatic C-H bending), 873.92 (N-N), 3208.00 (OH stretching), 1602.00 (Aromatic C=C stretching for B-ring and C-ring), 1353.00 (OH bending of phenol);^1^H NMR (DMSO, 500 MHz, ppm)-5.8(d),6.69(s), 6.51(s), 6.60(s),7.72(s),7.61(s),6.61(s), 7.23(s), -OH(5.0), -OH(15.0).

### QUESH

Colour-Orange; M.P.-366°C; Mol wt.-436.37 g/mol; t_R_-5.613 min; FT-IR (cm^−1^): 1754.00 (C=O), 1293.60 (Aromatic C–H), 783.95, 691.91, 570.60 (Aromatic C-H bending), 874.46 (N-N), 3199.00 (OH stretching), 1606.00 (Aromatic C=C stretching for B-ring and C-ring), 1354.00 (OH bending of phenol);^1^H NMR (DMSO, 500 MHz, ppm)-5.8(d),6.69(s), 6.51(s), 6.60(s), 7.78(s), 7.0(s), 7.34(s), 6.91(s).

### Computational Analysis

#### Drug Likeness and ADME Analysis of QDs

The screened derivatives were analyzed for ADME properties using the SwissADME web tool (http://www.swissadme.ch/) (Daina *et al*. 2017) This tool helps in predicting pharmacokinetic parameters such as blood-brain barrier (BBB), plasma protein binding (PPB), gastrointestinal (GI) absorption, Lipinski rule of five, solubility, total molecular polar surface area (TPSA), bioavailability, and drug score.

#### Molecular Docking of Synthesized QDs within Binding site of PPAR-γ

Molecular docking was performed using the AutoDock/Vina plugin with scripts (http://autodock.scripps.edu/) (Trott and Olson 2010) from the AutoDock 1.5.6 Tools package. The 3D crystal structure of PPAR-γ (PDB code: 4JAZ) was retrieved from Protein Data Bank (PDB) (https://www.rcsb.org/) bound with agonist trans-Resveratrol (resolution: 2.85Å). First, the receptor and ligand structures were transformed to the PDBQT file format, which includes atomic charges, atom-type definitions, and for ligands, topological information (rotatable bonds). The protein structure was prepared for the domain cavity by assigning polar hydrogen atoms with the subsequent addition of the Kollman and Gasteiger charges. A grid with x, y, and z dimensions of approx. 70 × 70 × 70 grid points was set up to cover the 3-dimensional active site of the PPAR-γ.

Afterward, ACD/ChemSketch was employed to draw the ligands (synthetic derivatives) and saved in MOL file format. While, the 3D structures of quercetin (CID:5280343) and RSG (CID:77999) were used as a reference compound from the PubChem database. The energy of each ligand was minimized in Avagadro (https://avogadro.cc/) using the Steepest Descent method and Merck molecular force field [MMFF94(s)] (Lim *et al*. 2020) to get energetically stable confirmation. Finally, the test and reference ligands were read into the AutoDock tool, and the best conformation was ascertained.

#### MD simulation Using Desmond

After the final refinement, QUESH attained the best interaction with PPAR-γ and hence, the QD was subjected to further Molecular Dynamics (MD) simulation along with reference compounds; quercetin and RSG using the Desmond package (Schrodinger release 2020). A timescale of 100ns was employed using the Optimized Potentials for Liquid Simulations-2005 (OPLS-2005) force field with a temperature of 300K and pressure of 1 bar with NPT (constant number of atoms N, pressure P, and temperature T) ensemble. The key parameters opted constitute the Nose Hoover chain thermostat method constraints, Martyna-Tobias-Klein barostat method, and integrator as MD for attaining stable equilibrium. The complex systems were subjected to solvation by implementing explicit water-filled cubic box buffered at 10 Å spacing with three-point water model (TIP3P) with periodic boundary conditions. Further neutralization of the net charge of all the complex systems was ensured by adding sodium ions (Na+).

### Biological Evaluation

#### Cell culture conditions

The human NSCLC cell lines A549 and NCI-H460 were procured from the National Centre for Cell Science (NCCS), Pune, Maharashtra, India. Both the cell lines were maintained in RPMI-1640 medium supplemented with 10% FBS, 1% 200 U/mL of penicillin, 1% 0.2 mg/ml of streptomycin, and 1% 200mM L-glutamine in a humidified, 5% CO_2_, 37°C incubator (Eppendorf, Germany). The media was changed every 2-3 days, and the cells were separated via trypsinization using 0.25% trypsin/0.038% EDTA when they reached 70-80 % confluence.

#### Cytotoxicity determination

The potential cytotoxic effects of PPAR-γ ligands (synthesized QDs) were examined using an MTT assay as per the standard protocol (Li *et al*. 2020; Paul *et al*. 2008). Briefly, 1 × 10^5^ cells were plated per well onto 96-well plates (Thermo Scientific, USA) and kept overnight to confluence. 24h post-plating, cells were treated with various concentrations of QDs and controls; quercetin, and RSG (0.1, 0.01, 0.001, and 0.0001 mM) in a growth medium for two timelines, 48 h and 72h. 20µl of the freshly prepared 3-(4,5-dimethylthiazol-2-yl)-2,5-diphenyl-2H-tetrazolium bromide (MTT) solution (5 mg/ml in PBS) was added to each well and incubated for 4 h (37°Cand 5% CO_2_). After incubation, the medium was aspirated, and the formazan crystals were dissolved by adding 200 µl of DMSO. The viability of cells was determined by measuring their absorbance at 570 nm and 690 nm in a microplate spectrophotometer (Epoch Microplate Spectrophotometer, Biotek). The optical density of absorbance is directly proportional to the number of live cells. The IC_5_, IC_10_, IC_20_, IC_30_, IC_40_ and, IC_50_ concentrations for all the drugs were calculated.

#### Scratch Assay

1×10^6^ cells (A549 and NCI-H460) were seeded into a 6-well plates (Thermo Scientific (USA) and after 24h post-plating, they were given serum starvation for the next 24h. The next day medium was aspirated, and the monolayer was wounded by manual scratching of a straight line with the help of a sterile 10μl micropipette tip to create a gap of constant width. Subsequently, cellular debris was washed with PBS, and the cells were then exposed to QDs and reference compounds (IC_5_ dose) with serum-free medium for 30h under standard conditions. The wound closure was monitored, and images were captured at 0 h, 24 h, and 30 h, respectively, with the help of Magnus Analytical MagVision, Version: x64 equipped with a camera (Magnum DC5) whereby the gap distances of the wounds were measured using the following formula (Bobadilla *et al*. 2019):

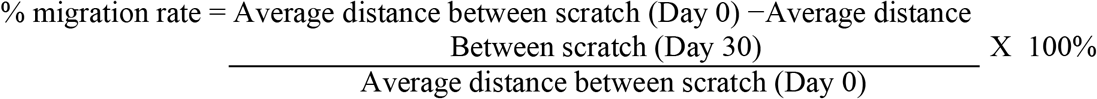

#### Western blot analysis

A549 cells were treated with IC_5_ doses of QDs and positive controls, quercetin and RSG for 24hr and thereafter cell lysates were prepared. Protein concentrations of all the mentioned samples were determined using the Bradford assay. An equal amount of protein was boiled at 95°C for 5 min after the addition of 3X Laemmli buffer, followed by separation on 10% SDS-PAGE and the separated proteins were electrotransferred onto polyvinylidene fluoride (PVDF) membrane using a Trans-Blot® Turbo™ Transfer System (Bio-Rad Laboratories, Inc., India). The membranes were first blocked with 5% non-fat milk in TBST for 1h at room temperature, followed by probing with respective primary antibodies following published protocols (Li *et al*. 2020; Paul *et al*. 2014). The primary antibodies used are as follows; anti-PPAR-γ (1:500), EMT markers-anti-Snail (1:500), anti-N-cadherin (1:500), anti-Vimentin (1:1000), anti-Slug (1:250), anti-E-cadherin (1:500) and anti-Zeb-1 (1:250). Anti-rabbit antibody conjugated to horseradish peroxidase (HRP) (1:1000) was used as the secondary antibody and incubated for 1h. β-actin was used as a control. The blots were incubated in ECL Plus reagent, and chemiluminescence was detected on iBright CL1500 Imaging System (Thermo Fischer Scientific) and quantified the density of the bands.

#### Cell cycle assay

2.5 × 10^5^ cells/well (A549 and NCI-H460) were seeded in 6-well plates and after 24h post-plating, they were treated with all the compounds (IC_5_ dose) for 48h. After incubation, the cells were rinsed with PBS, trypsinized, and collected by centrifugation. The cells were further washed and fixed overnight in 70% ethanol at 4°C. The pellet obtained was resuspended in a staining solution containing 50 µg/ml propidium iodide and 100 µg/ml RNase A for 30 min. The suspensions were then analyzed using a Becton Dickinson FACS Calibur™ system (BD Biosciences, Franklin Lakes, NJ, USA).The percentage of cells was determined in the G0/G1, S, and G2/M phases of the cell cycle by their DNA contents and presented as a fold of control (Ravichandran *et al*. 2014).

#### Statistical analysis

Normally distributed continuous variables are expressed as the means ± standard deviation (SD). One-way Analysis of Variance (ANOVA) followed by the “Dunnett’s Multiple Comparison Test” method was used to compare the results obtained in different treatment groups. A two-sided P value < 0.05 was considered statistically significant. Statistical analyses were performed using GraphPad Prism version 5.0 software (GraphPad Software, La Jolla, CA, USA).

## RESULTS AND DISCUSSION

### Chemistry

It is observed that the carbonyl group (>C=O) of ring C of quercetin aids a greater interaction when substituted with Schiff base (Ravichandran *et al*. 2014).This structural modification is reported to increase the water-solubility and stability of quercetin. Figure 1 illustrates the typical reactions involved in the formation of Schiff base derivatized compounds; TSC and hydrazones. The schiff-based quercetin complexes were synthesized by treating an appropriate amount of quercetin with the specific derivatives of quercetin, following their respective synthesis route outlined in Scheme 1. The process yielded colour products (QUETSC-yellow; QUEINH-light yellow; QUENH-light yellow; QUE2FH-orange; QUESH-orange) ranging from 65% to 80% after recrystallization with absolute ethanol, which were solid at high M.P. of 370°C, stable in air and soluble in an organic solvent, DMSO. All the spectras obtained from chromatographic and spectroscopic techniques have been provided in the supporting information (Figure S1-S3).

**Fig. 1.**
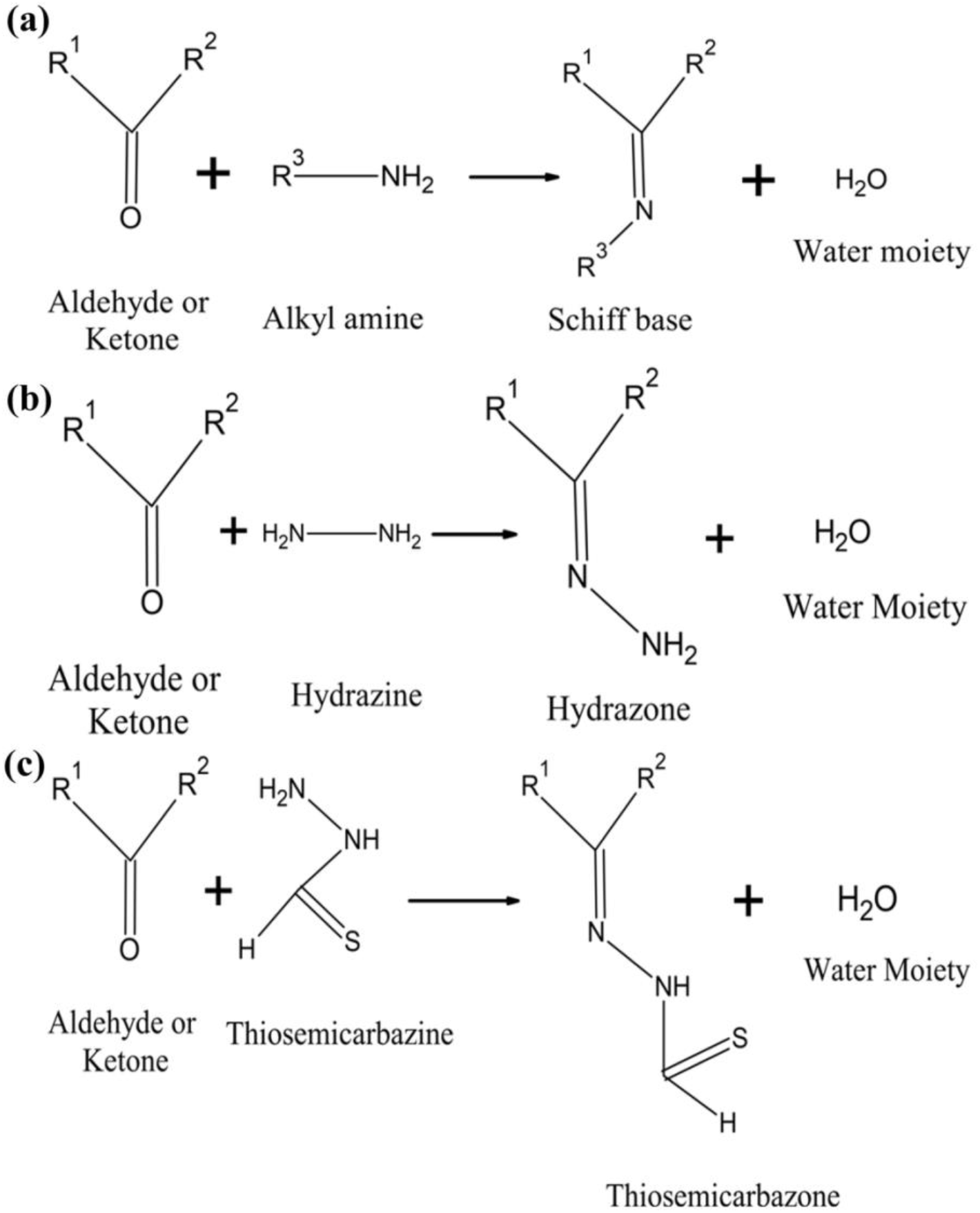
Typical reaction route of Schiff base and their derivatives. (a) Formation of Schiff base, (b) Formation of TSC, and (c) Formation of hydrazone.

The progress of each reaction was monitored through TLC using chloroform: methanol (9:1) solution as the mobile phase. HPLC was performed to affirm the purity of the synthesized compounds where the peaks were well eluted at different retention times (Figure S1). The resultant peaks are in full agreement, suggesting that the analysis was selective to the medium and excipients showed no interference with any excipient peaks. The effect of ligand-binding event was mapped using FT-IR analysis (Figure S2) where all the peaks were visible originating from the drugs. Spectroscopic technique, ^1^H NMR, was run to deduce the structural information of synthesized schiff base compounds and examine the differences in chemical shift (Figure S3). It was conferred that based on the chemical shift of QDs, the resonance at 7.0ppm (d) attributes to the azomethine group due to the involvement of amide (-NH-C=O) proton except for QUETSC, where the –NH proton groups with sulfur (-NH-C=O). The pyridine ring of QUEINH resonates between δ 7.96 to 9.06 ppm. This resonance was observed to be slightly different in compound QUENH where the nitrogen atom is situated at the meta position, falling between δ 7.63 to 9.17 ppm (Figure S3 (c)). The furan structure of QUE2FH displays a signal between δ 6.60 to 7.72 ppm (Figure S3 (d)), which corresponds to >C=O while the benzene ring of QUESH resonates between δ 6.91 to 7.78 ppm (Figure S3 (e)), ensuring the existence of aromatic protons. Hence, the ^1^H NMR data indicated that the structure modification widely supports reaction with the imine nitrogen replacing the carbonyl group (C=O) of the C ring (Chen *et al*. 2011).

**Scheme 1:**
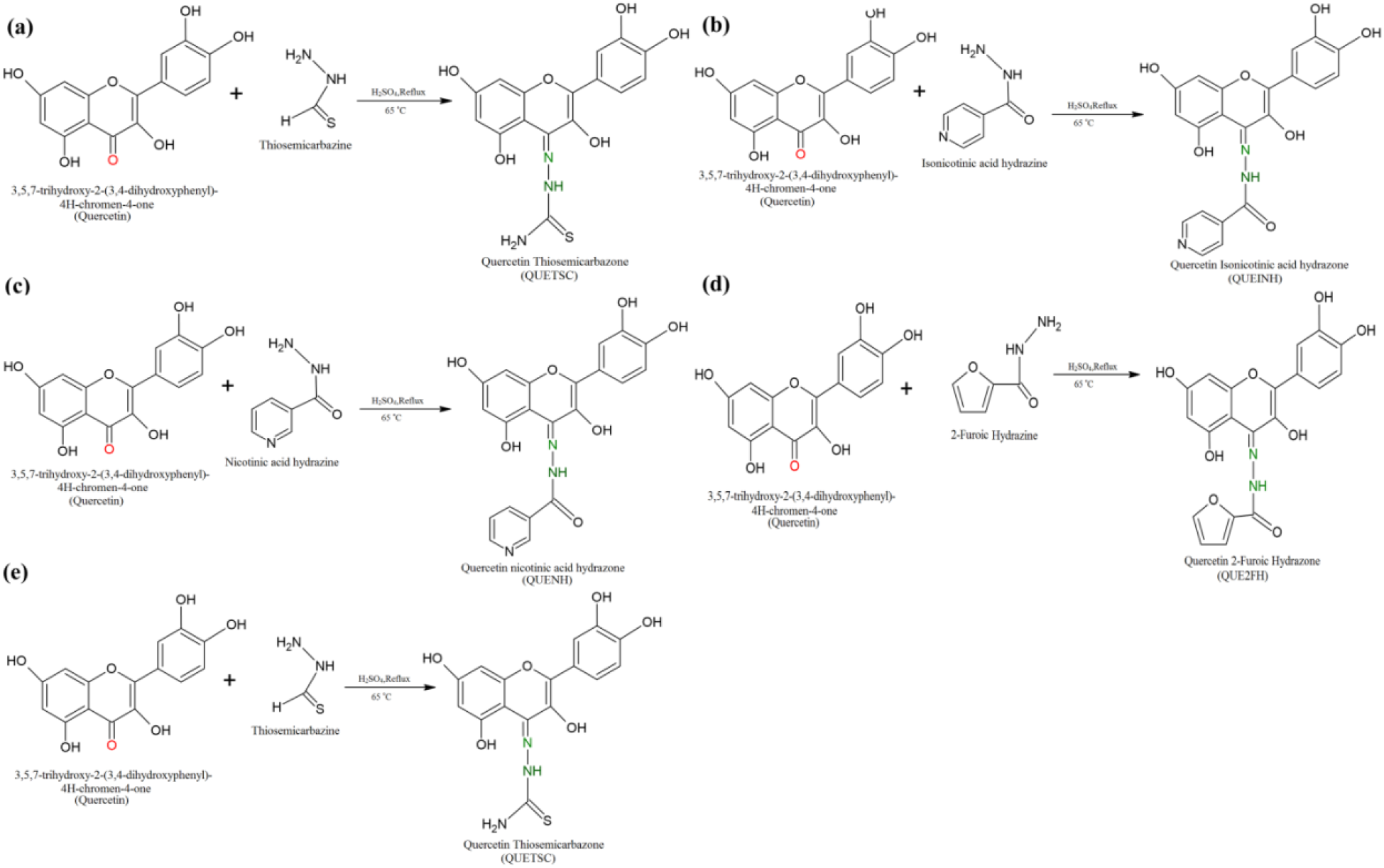
Synthetic route of QDs. (a) QUETSC; (b)QUEINH; (c)QUENH; (d)QUE2FH (e)QUESH

### In silico ADME Prediction profiles of QDs

SwissADME (Daina *et al*. 2017) web tool encompasses a user-friendly interface that pools out good ADME profiles of predictive models and generates the report in the form of a Bioavailability Radar that allows a first glance at the drug-likeness of a molecule. Figure 2 represents the bioavailability radar of the synthesized QDs, wherein the pink area represents the optimal range for each property (Chen *et al*. 2016). As noticed, all compounds fall entirely in the pink area (except for unsaturation fraction), affirming their efficacy of possessing superior drug-like properties.The obtained TPSA values, as tabulated in Table 1,ranged from 164 to 196 Å^2^ which engrosses the violation of lipophilicity and hence prompts a factor of reduced gastrointestinal(GI) absorptive level. However, the overall result attests that the absorptivity level of all the QDs has been improved as the Estimated SOLubility (ESOL) Class is found to be better than quercetin. Another pharmacokinetic property was observed that they all are not permeability glycoprotein (P-gp) substrates, which makes them a successful drug against multidrug-resistant cancer cells, overexpressing this drug transporter. Other drug distribution descriptors are projected in supporting information (Table S1-S2).

**Table 1.**
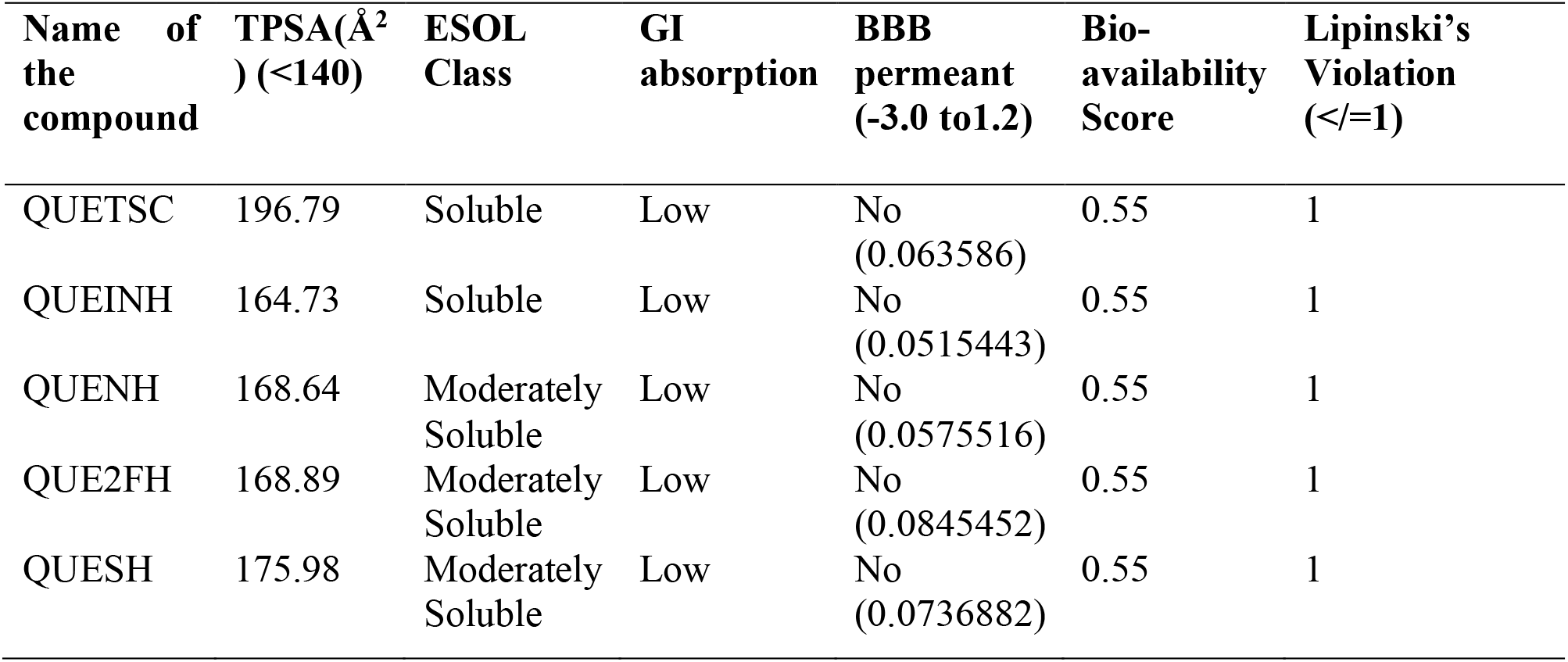
ADME properties of synthesized QDs using SwissADME web tool.

**Fig. 2.**
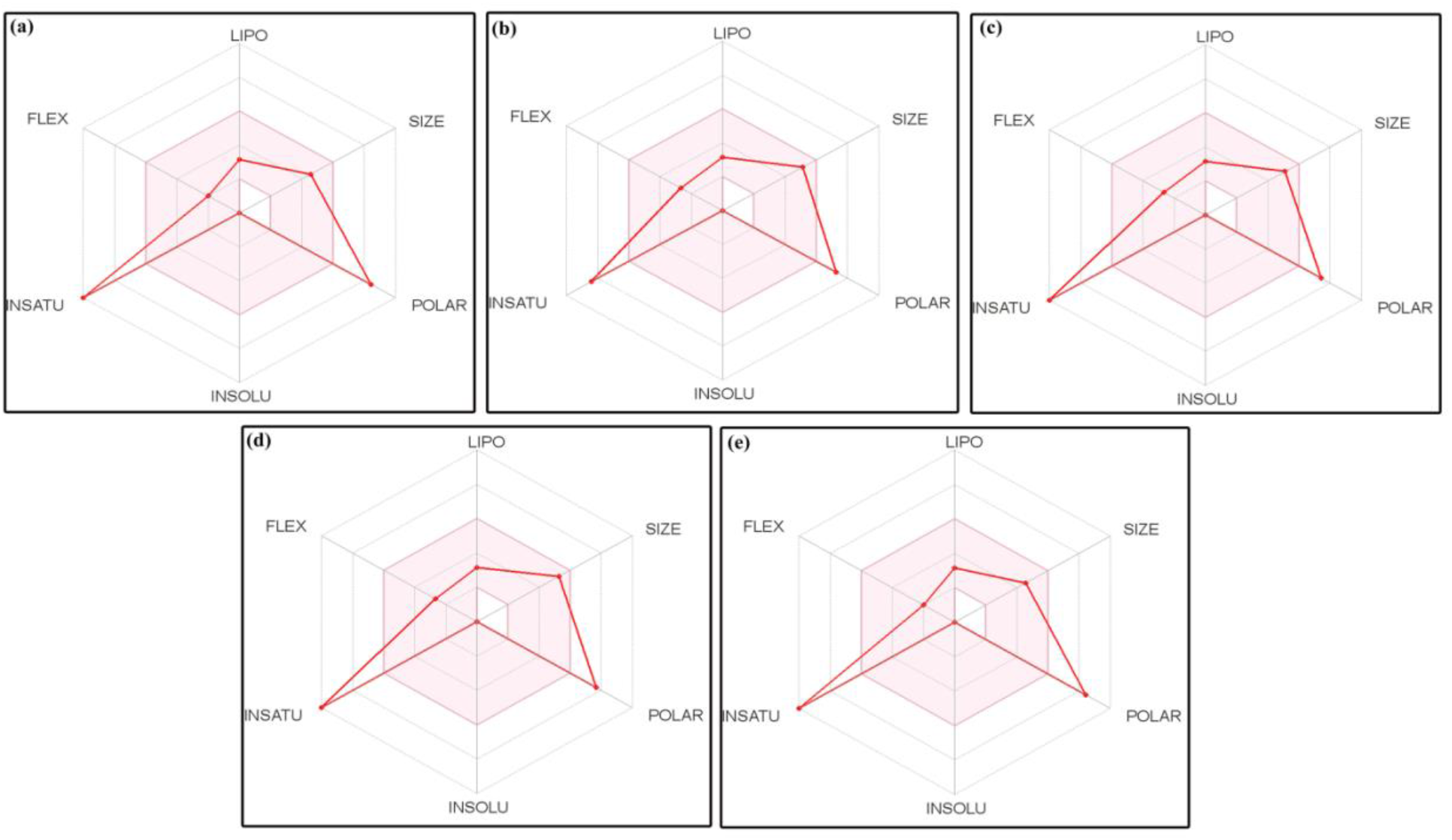
Bioavailability radar of QDs based on their suitable physicochemical indices ideal for oral bioavailability using SwissADME predictor; (a)QUETSC; (b) QUEINH; (c)QUENH; (d)QUE2FH (e) QUESH (Lipophilicity (LIPO): XLOGP3 between -0.7 and +5.0, Molecular weight (SIZE): MW between 150 and 500 g/mol, Polarity (POLAR) TPSA between 20 and 130 Å 2, Solubility (INSOLU): log S not higher than 6, Saturation (INSATU): fraction of carbons in the sp3 hybridization not less than 0.25, and Flexibility (FLEX): no more than 9 rotatable bonds.)

### Binding Interaction Profile of PPAR-γ Ligands with its Receptor

In order to assess the question of whether PPAR-γ is specific towards structural modifications, we employed computational analysis involving molecular docking and MD simulation techniques. According to the docking energy framework through Autodock, all the ligands; QUETSC, QUEINH, QUENH, QUE2FH, and QUESH were well accommodated in the PPAR-γ binding site and accorded a significant improvement (Figure 3). Upon the associated physical-chemical molecular interactions, the docking scores with interaction residues, bond type, and bond distance are summarized in Table 2. QUEINH and QUESH exerted greater stability with the receptor amongst the five derivatives with a docking score of -10.1 and -10.5 kcal/mol, respectively. This might be due to the formation of Hydrogen-bonds (H-bonds) as QUEINH attained a total of six interactions, while QUESH formed 10 interactions with amino acid residues. It is well known that H-bonds form the strongest bonds and are conserved in most agonists of PPAR-γ, which plays a key parameter for its activity (Salmaso and Moro 2018).In order to compare the results, we used a positive control, rosiglitazone (RSG) which is the potent synthetic agonist for PPAR-γ. Conversely, we observed RSG to have a low binding energy value of -7.6 kcal/mol with the least H-bond. While quercetin was sighted to fit better than RSG in the binding pocket of PPAR-γ. Overall docking results revealed that the Schiff base compounds could effectively form pivotal contacts with the PPAR-γ. Particularly, QUESH was observed to occupy this pocket efficiently which highlights its increased flexibility.

**Table 2.**
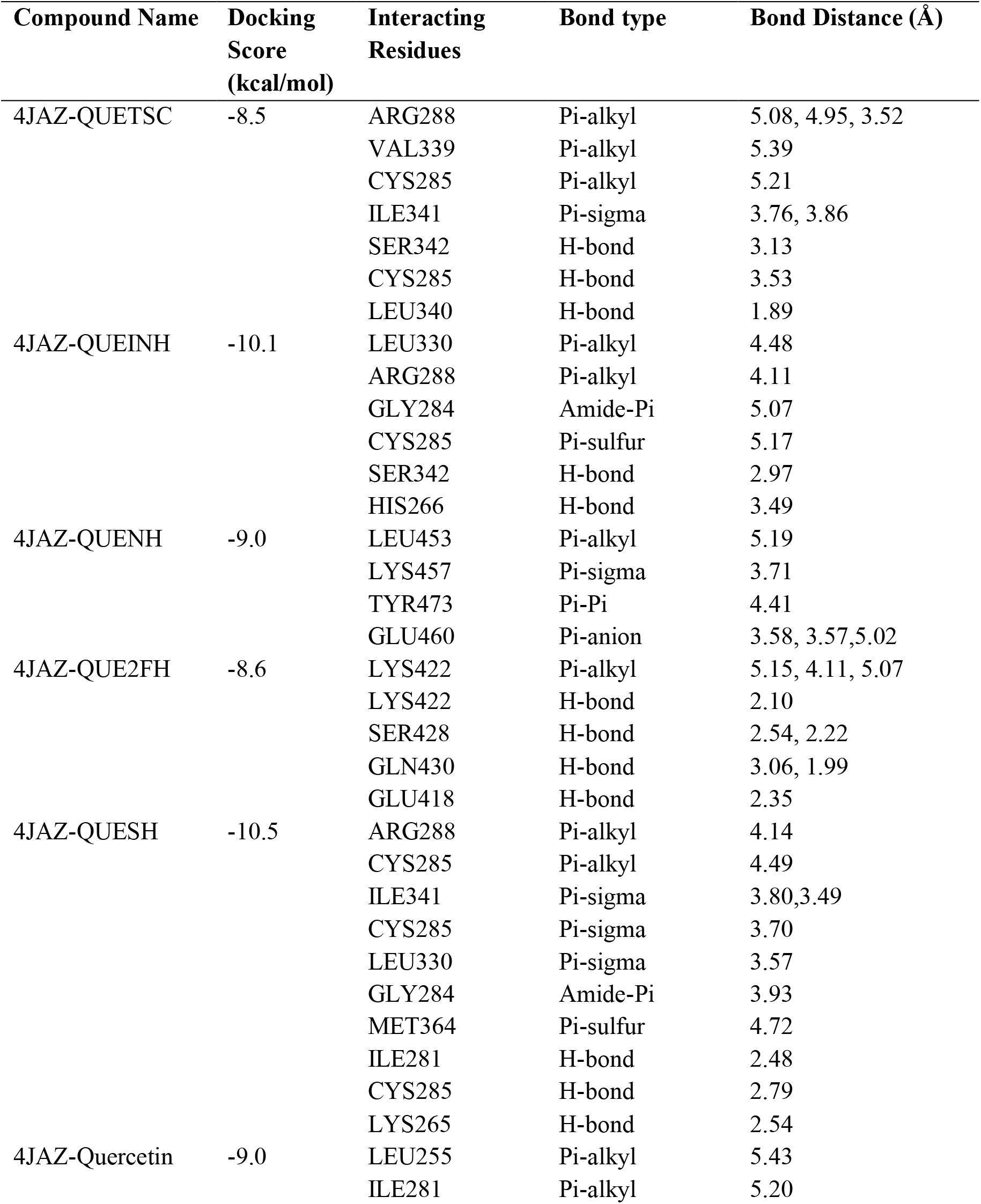
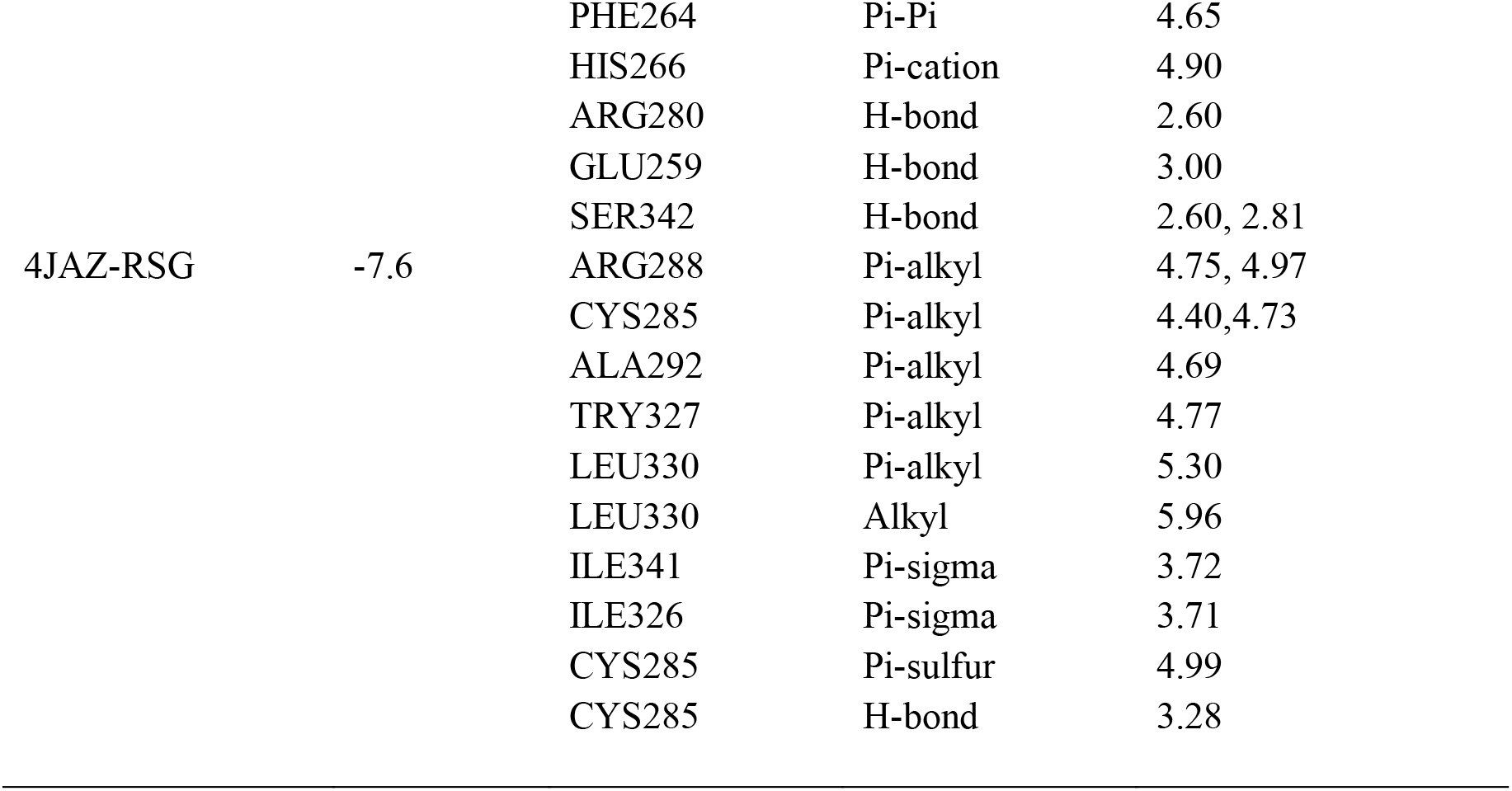
Interacting analysis of PPAR-γ Ligands with its receptor.

**Fig. 3.**
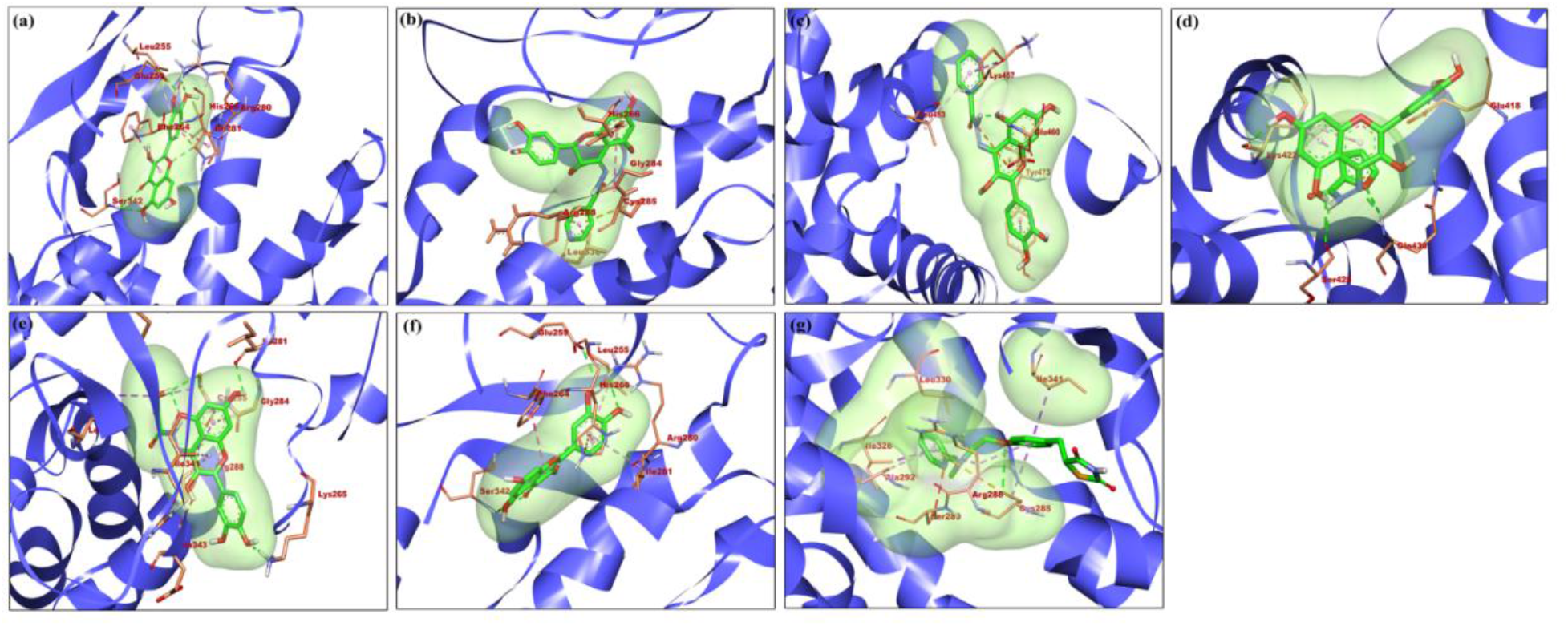
Interacting analysis of QDs with PPAR-γ receptor. Identified molecular docking interaction profile of (a) 4JAZ-QUETSC (b) 4JAZ-QUEINH (c) 4JAZ-QUENH (d) 4JAZ-QUE2FH (e) 4JAZ-QUESH (f) 4JAZ-Quercetin (g) 4JAZ-RSG acquired within the binding cavity of PPAR-γ-LBD in 3D format generated through Discovery Studio 4.1 Visualizer.

### MD Simulation: Structure Basis Relationship

MD simulation is considered as the most reliable computational tool for investigating the structural relationship between a drug and a protein. Therefore, we followed the Desmond MD protocol with QUESH due to its increased flexibility towards the PPAR-γ receptor target. Table 3 specifies the prominent parameters required to stabilize the system. As revealed by the graphical plots of Figure 4, the stability of 4JAZ-QUESH was consistent and stabilized during the entire 100ns process of MD simulation. In PPAR-γ, QUESH maintained for 95% of the total simulation time as compared to RSG with 78% and quercetin with 60%. Generally, it has been speculated that a protein structure comprising fewer deviations aids in flexible conformation irrespective of turns, bends, and coils involved, which are truly accomplished by lower RMSD and RMSF values (Pappa *et al*. 2007).On monitoring the simulation trajectories, it was inferred that 4JAZ-QUESH recorded the least deviation with a constant RMSD value of 2.3Å and caused relatively low RMS fluctuations (Figure 4 (A & B)). Furthermore, protein-ligand contacts implied that 4JAZ-QUESH complex established a larger amount of H-bonds and hydrophobic contacts, which is in agreement with the above docking results and elucidates it to be a good candidate for PPAR-γ. Merely, the presence of weak intermolecular interactions (H-bonds and hydrophobic interactions) in 4JAZ-QUESH addresses the overall stabilization of the complex over the reference compounds (Salmaso and Moro 2018).

**Table 3.**
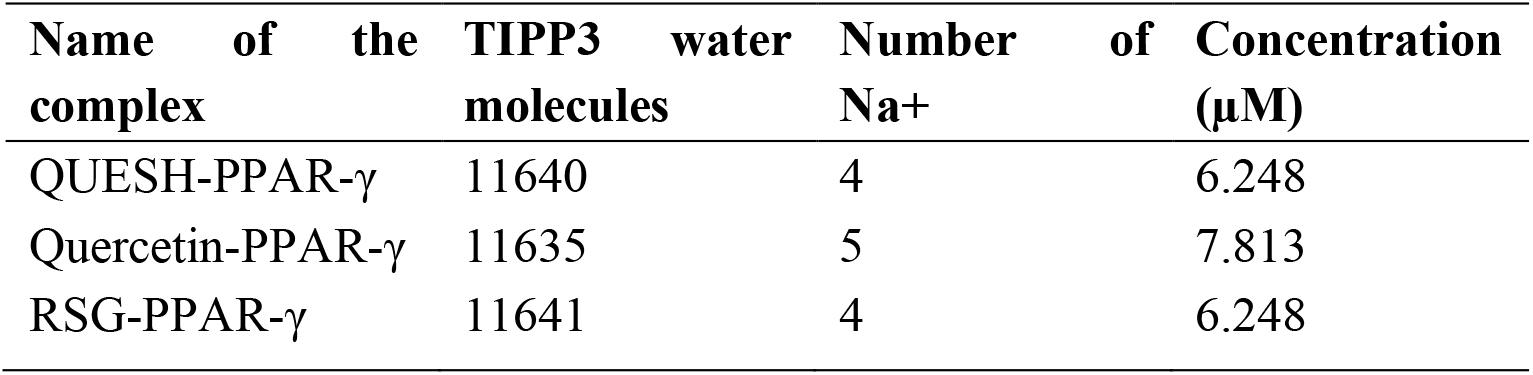
Governing parameters required for stabilization of the complex system for MD simulation.

**Fig. 4.**
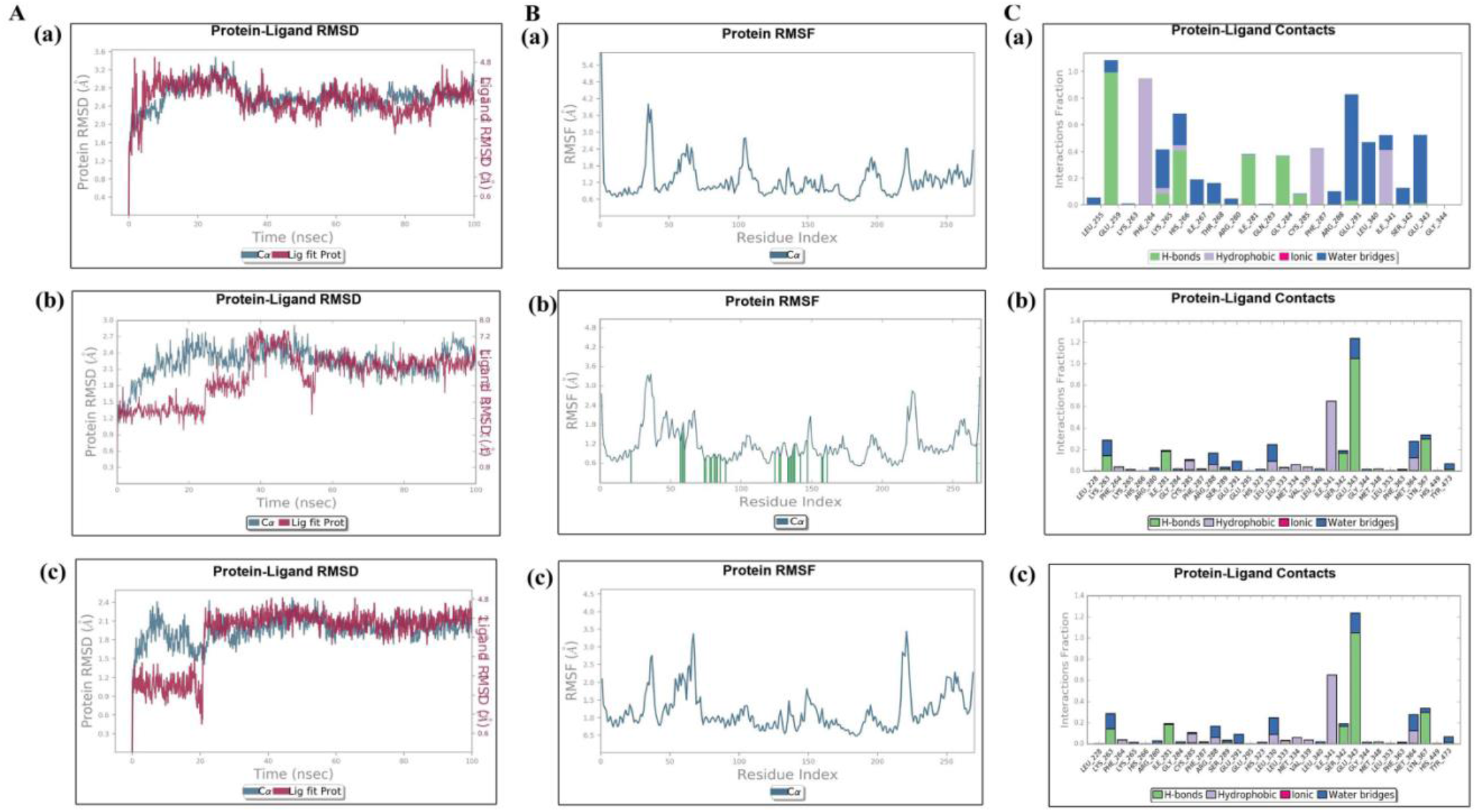
MD simulation of PPAR-γ partial agonist. The calculated RMSD values (A), RMSF values (B), and protein-ligand contacts (C) of (a)-QUESH and reference ligands; (b)-Quercetin and (c)-RSG with PPAR-γ.

### PPAR-γ Based Ligands-Synthesized QDs reduced cell viability of A549 and NCI-H460 cells

Next, we tend to investigate the activity of synthesized QDs on the proliferation rate of A549 and NCI-H460 cells through MTT assay. Table 4 summarizes the IC_50_ values obtained for all the drugs in A549 and NCI-H460 cells. QDs has ascribed dose and time dependent anti-proliferative effects on both the lung cancer cell lines leading to excellent and estimable inhibitory concentration values. It is clear from the graphs that RSG unveiled a very mild effect on cell viability, while quercetin elicited a better outcome than RSG (Figure 5). Among all the synthesized QDs, QUETSC and QUESH halted cell growth of A549 to 13-fold at 72h post-treatment. They reduced the survival rate of A549 cells and attained approximate maximal effects at 1.03±0.33µM and 1.11±0.05µM, respectively, compared to NCI-H460 cells which showed no noticeable change. This significant characteristic feature of both the QDs was due to the presence of electron-withdrawing group (QUETSC; >C=S and QUESH; >C=O) that results in further active delocalization of nitrogen lone pair of azomethine groups of the compounds (Pappa *et al*. 2007).As QUETSC and QUESH attributed greater cytotoxicity, thereafter, we sought to mark out the effects on cell cycle distribution of A549 and NCI-H460 cells. Interestingly, the proliferation of both the cells was not affected by the derivatives as compared to untreated control (Figure S4). The cells have reached complete G1-S with subsequent decrease in the percentage of cells dramatically in S and G2-M phase. Unfortunately, the incorporation of QDs reflected no arrest of the cells, suggesting that the two synthesized QDs are not responsible for apoptosis.

**Table 4.**
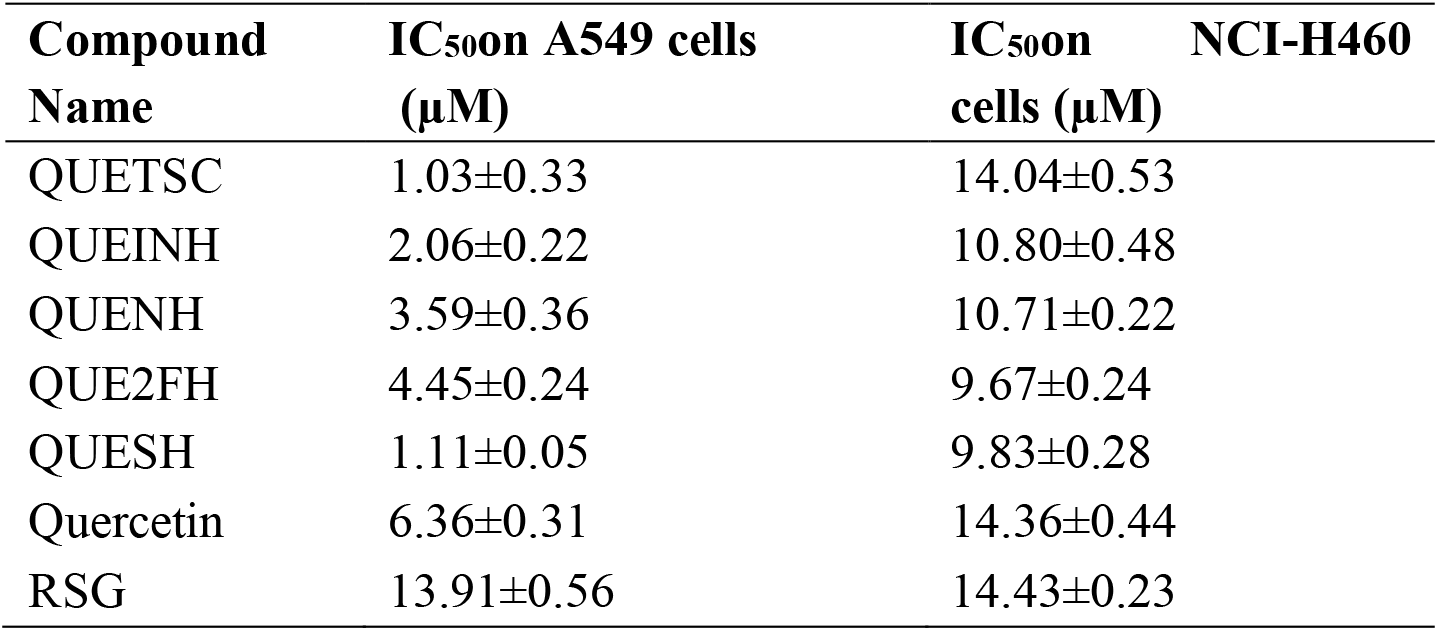
IC_50_ values of QDs with controls performed by MTT assay on A549and NCI-H460 cells.

**Fig. 5.**
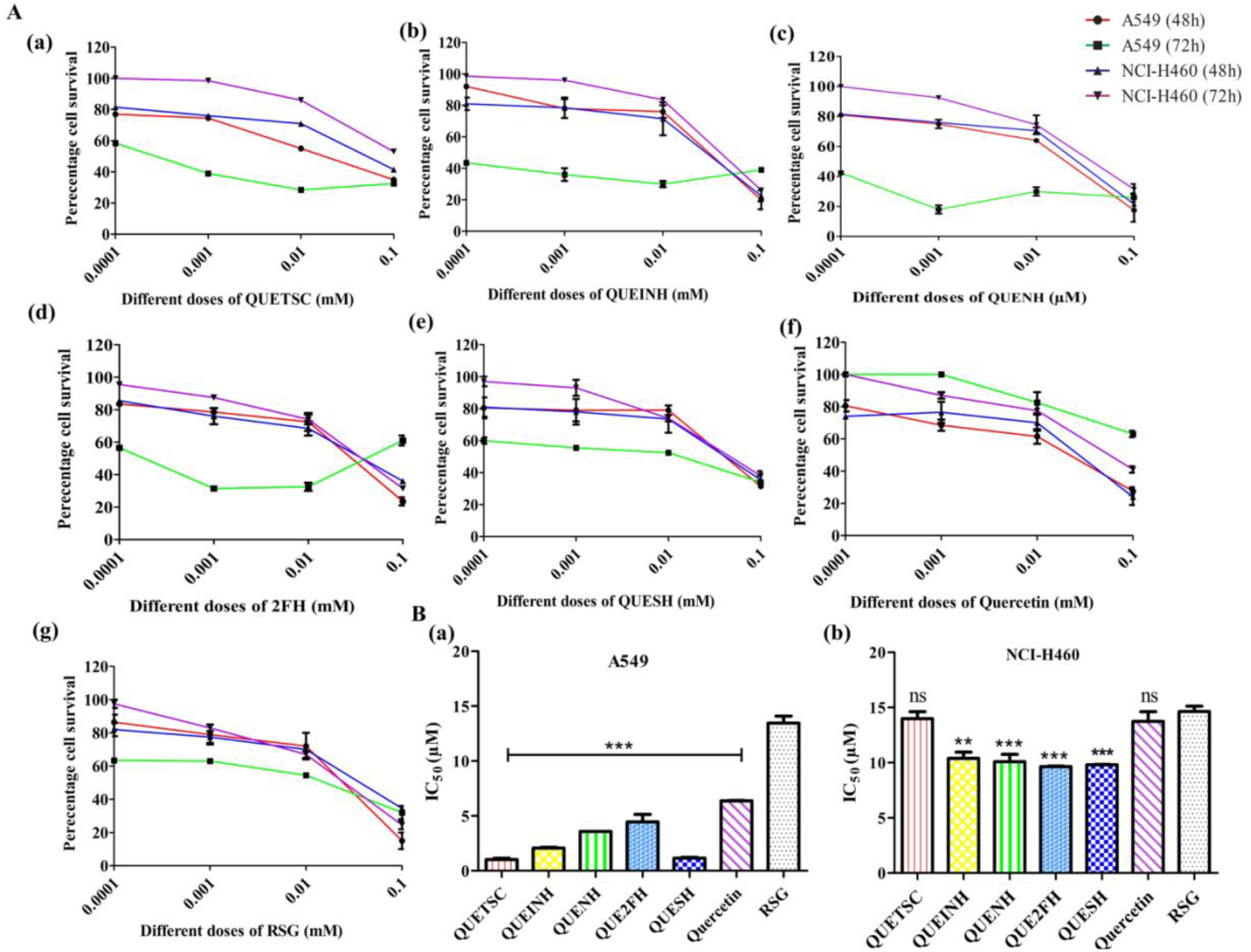
Anti-proliferative effects of PPAR-γ based ligands (synthesized QDs) on A549 and NCI-H460 cells. The cell viability was determined by treating cells with the compounds of varied concentrations (0.1, 0.01, 0.001 and 0.0001 mM) for 48h and 72h using the MTT assay. (A) Survival ratio of A549 cells and NCI-H460 cells at two different time intervals by QDs-(a) QUETSC; (b) QUEINH; (c) QUENH; (d) QUE2FH; (e) QUESH; (f) Quercetin and (g) RSG. (B) Representation of the half-maximal IC_50_ values of A549 cells (a) and NCI-H460 cells (b). Each value is presented as mean±SD for three replicate determinations for each treatment (p<0.05, one-way ANOVA). The error bar represents SD; *p<0.05, **p<0.01 and ***p < 0.001 compared with control group-RSG.

However, it is evident that the A549 cells are more sensitive than NCI-H460 cells (Figure 5(B)) as the cells exhibited minimal cytotoxicity toward QDs. The differential cell survival between the two NSCLC cells may be due to the higher proliferation rate of NCI-H460 cells than that of A549 cells, which promotes greater vascularity and the ability to metastasize (Chen *et al*. 2011; Manley and Waxman 2014).Reports suggest that a sub-population of NCI-H460 cells possess stem-cell-like property, which can self-renew with greater proliferative potential and enhanced tumorigenicity (Shi *et al*. 2012). Eventually, this increases drug efflux through trans-membrane drug transport proteins, leading to decreased intracellular drug accumulation. Hence, the cells might become resistant to any chemotherapeutic drug. This has been effectively shown by Pesic M. and coworkers, who reported the emergence of a novel cell line, NCI-H460/R, after doxorubicin treatment to NCI-H460 cells, which developed up to 96.2-fold resistance to the drug (Pesic *et al*. 2006).The failure to sensitize the cells was attributable to an inability to prevent doxorubicin efflux. Based on these evidences, our results point toward a drug-resistant nature of the cell line NCI-H460.Hence, the effectiveness of tested compounds against A549 cell growth in descending order is: QUETSC>QUESH>QUEINH>QUENH>QUE2FH>Quercetin>RSG, whereas against NCI-H460 cells is: QUE2FH>QUESH>QUENH>QUEINH>QUETSC>Quercetin>RSG. Generally, it is considered that lower doses of drugs often signify to impart beneficial effect. Although for drug lead optimization, IC_50_ value is the most commonly preferred metric for on-target activity. However, there are studies where minimized doses of IC_50_ have been reported to be effective i.e., the lesser the value of IC_50_ the more potent is the drug (Meyer*et al*. 2019; Kalliokoski *et al*. 2013).Therefore, we selected nanomolar concentration i.e., IC_5_value of QDs for the subsequent mechanistic experiments. All the QDs induced higher toxicity in both NSCLC cells tested, which suggested therapeutically achievable efficacy.

### Exposure of PPAR-γ Based Ligands-Synthesized QDs inhibits A549 cell migration efficiently than NCI-H460

Tumor migration is well known for promoting metastasis. Following the trail of anti-proliferative effects, the migratory ability was examined after the treatment of QDs. We used a wound-healing assay to evaluate the migratory potential of the two NSCLC cell lines. As shown in Figure 6, regardless of the difference in migrating ability, distinct wound closure of each cell line was monitored for 30 h. Table 5 summarizes the % migration rate of A549 and NCI-H460 cells for all the synthesized QDs. The closure of the gap was greatly inhibited by QUETSC and QUE2FH on A549 cells with % migration rates of about 37.09%±0.54% and 29.67%±0.46%, respectively at 30h post-treatment. Similarly, QUESH ought to act hold of migration with% migration rate of 37.63%±0.37%. The addition of the other two QDs, QUEINH and QUENH did not narrow the scratch wound of the cells. In contrast, negligible differences in the migrating capacity of NCI-H460 cells were observed for all the QDs and instead showed an increase in their motility compared to the untreated ones (Figure S5). This might be due to a higher growing capacity of NCI-H460 cells, as their growth rate is double that of A549 cells (Gomez-Casal *et al*. 2013; Kumar *et al*. 2013).These findings proved the potency of QUETSC, QUE2FH, and QUESH in preventing autonomous migration of A549 cells with limited sensitivity to NCI-H460 cells.

**Table 5.**
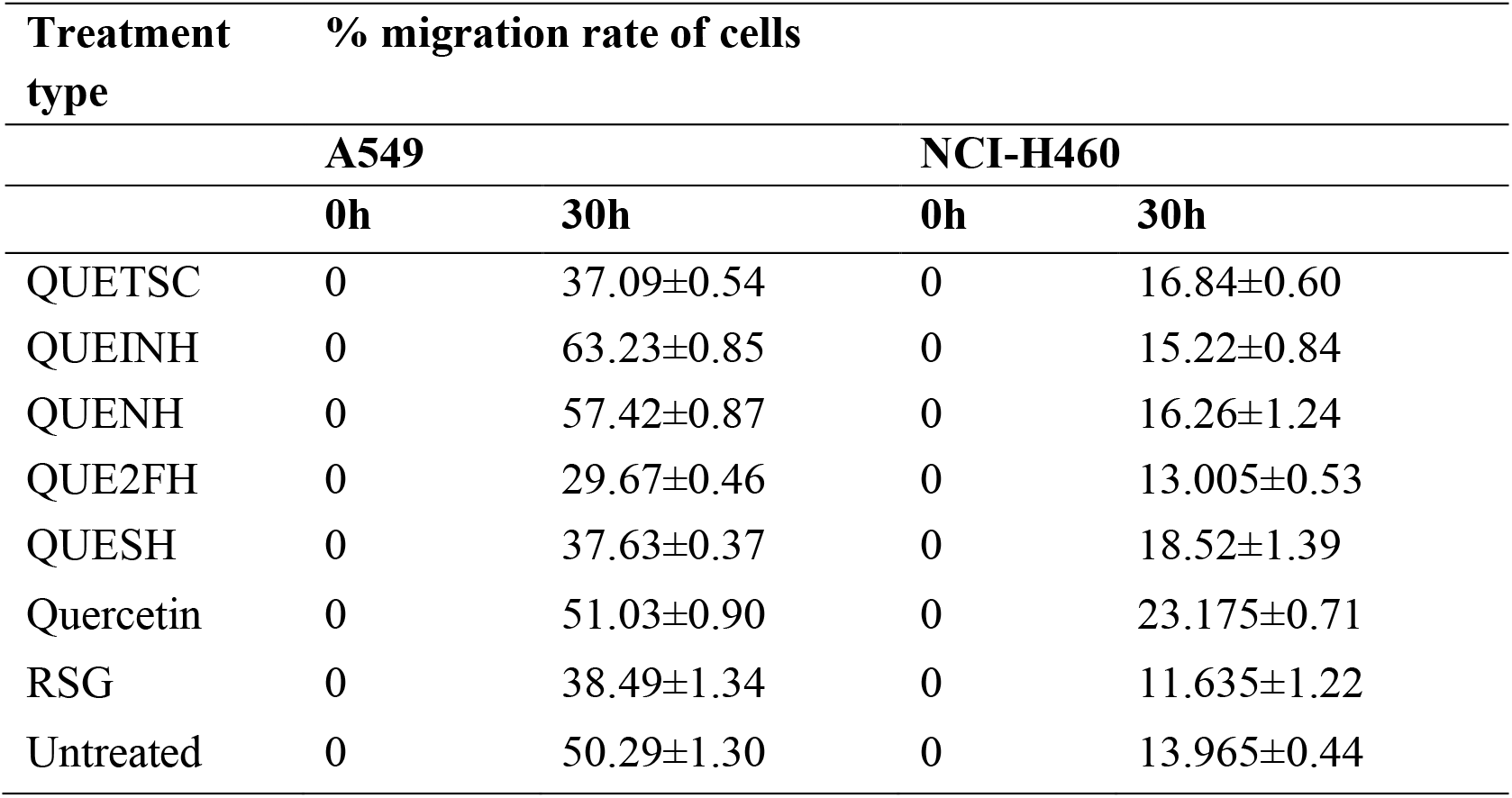
Percentage migration rate of A549 and NCI-H460 cells for PPAR-γ Based Ligands-Synthesized QDs.

**Fig. 6.**
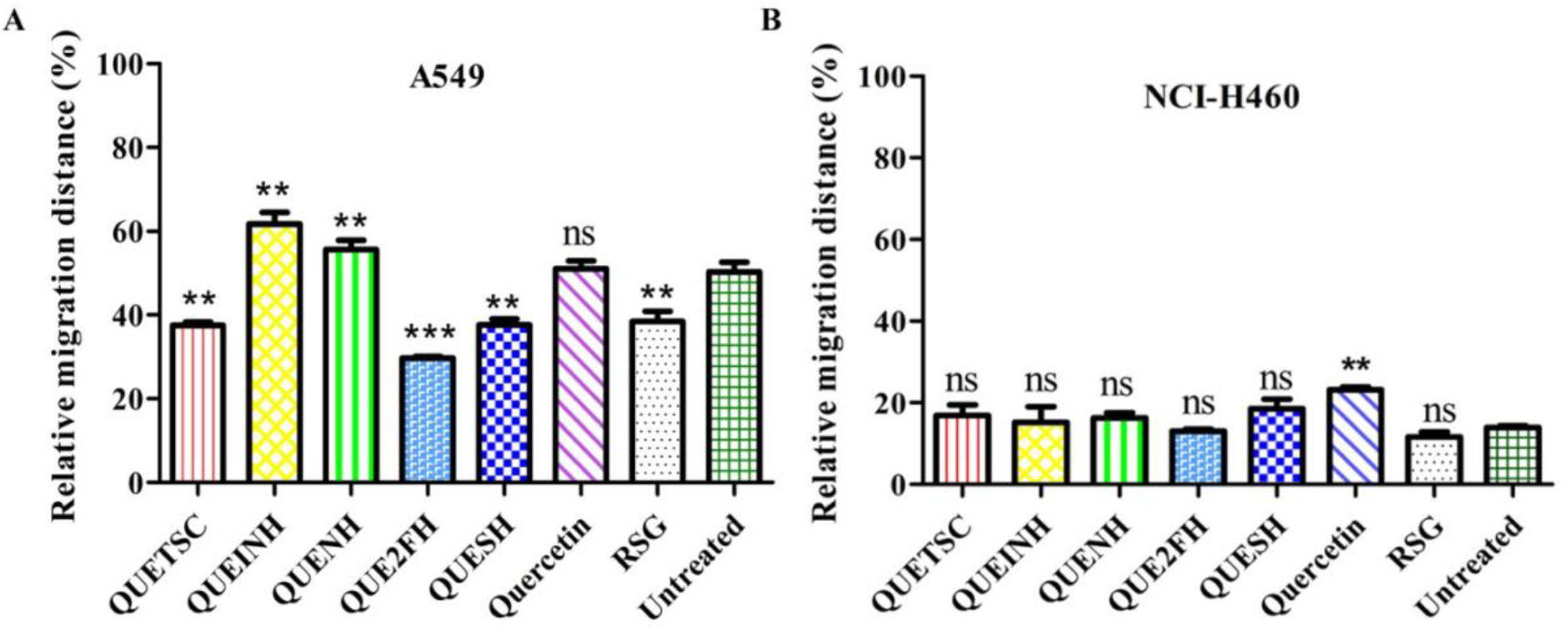
PPAR-γ Based Ligands-Synthesized QDs inhibit migration of A549 and not in NCI-H460 cells. The wound closure was quantified by measuring the remaining cell-free gap area under an Olympus microscope equipped with a camera (Magnus Analytical MagVision, Version: x86). Each value is presented as mean±SD for three replicate determinations for each treatment (p<0.05, one-way ANOVA).The error bar represents SD; *p<0.05, **p<0.01 and ***p < 0.001 compared with control group-Untreated cells.

### Expression of PPAR-γ protein in the two NSCLC cell lines

As QDs modestly retarded cell growth in A549 cells, we further investigated their effects on the activation pattern of PPAR-γ. The expression of PPAR-γ was primarily evaluated on both the NSCLC cells by employing Western blot analysis. The relative expression of PPAR-γ has been reported in Figure 7 (A and B), which confers the molecular mass of PPAR-γ protein to be 56 kDa. In A549 cells, PPAR-γ was expressed upto15-fold higher as compared to NCI-H460 cells which underscores that the amount of PPAR-γ protein varies in different cell lines. One reason might be due to the binding profile of PPAR-γ that are highly specific to the sites which avoid other repressive mechanisms and tend to sensitize a variety of cancer cell types, thereafter, subjecting to be cell type-specific (Nielsen *et al*. 2006).Thus, A549 cells are more specific towards PPAR-γ and hence chosen for subsequent analyses.

**Fig. 7.**
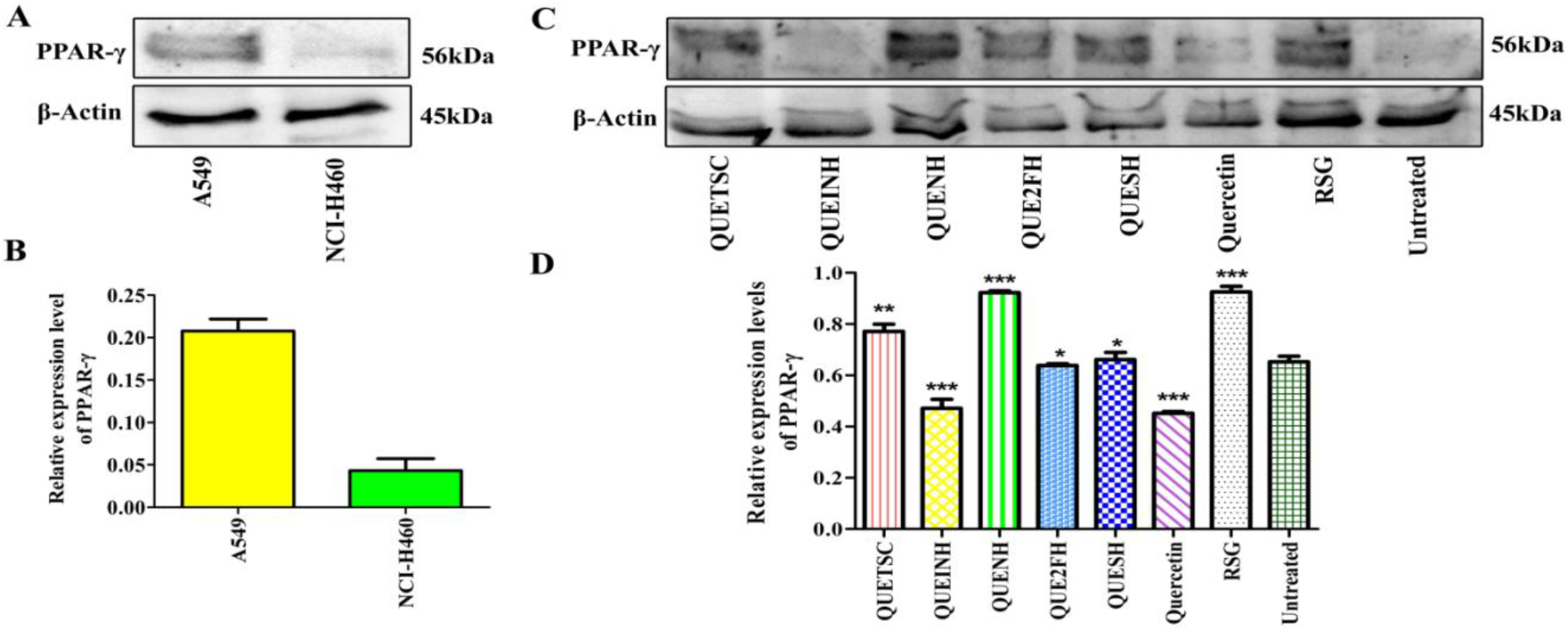
Synthesized QDs successfully activated PPAR-γ. (A & B) Expression and quantification of PPAR-γ at the protein level in A549 and NCI-H460 cells. (C & D) Expression and quantification of PPAR-γ at the protein level in A549 by synthesized QDs after 24h treatment. Each value is presented as mean±SD for three replicate determinations for each treatment (p<0.05, one-way ANOVA). The error bar represents SD; *p<0.05, **p<0.01 and ***p < 0.001 compared with control group-Untreated cells.

### QDs triggered PPAR-γ in A549 cells at the protein level in a ligand-dependent manner

Due to pronounced differences between the responses of A549 and NCI-H460 cells towards PPAR-γ, further examination of synthesized QDs on PPAR-γ activation was assessed in A549 cells. Figure 7 (C) connotes all the new synthesized ligands functioned as PPAR-γ activators. The speculated reason behind this activation is a large ligand-binding cavity of PPAR-γ that consists of polar residues at both of its ends and allows them to link with a wide array of ligands (Nielsen *et al*. 2006).It is worth noting that QUETSC, QUE2FH and QUESH displayed partial activation of PPAR-γ with a significant lower intrinsic activity. As mentioned, polar residues is thought to be mediate the full/weak/partial activation of PPAR-γ. The presence of amino acids associated with PPAR-γ binding domain involves Tyrosine, Histidine, Serine, Isoleucine and Cysteine (Shang and Kojetin 2021). With the interacting residues hosted by QUETSC, QUE2FH and QUESH seemed to follow a distinct differential pattern of stabilizing the regions of ligand binding pocket (Figure 3 (a, d, f)). The combination of CYS285, ILE341, ILE281, SER342 and SER428 amino acid residues are majorly responsible to reinforce this sort of stabilization which determines the depth of structural feature.

However, QUEINH manifested lowest potency on PPAR-γ that led to similar level of expression to that of untreated cells. This reflects the nature of QUEINH as weak agonist. The highest PPAR-γ expression was witnessed by QUENH that is comparable to that of RSG and describes it’s a full agonist. As expected, RSG displayed a higher level of PPAR-γ stimulation of about 40% increased expression with respect to untreated. This is because full agonist tends to interact directly with α-helix due to the presence of acidic heads (Yang *et al*. 2013). We can figure out from the docking picture of 4JAZ_RSG that the presence of TYR327 residue is thought to elicit full agonist efficacy. Similarly, QUENH involved TYR473 residue and also seen to directly bind to LBD region which is why it is correlating with full agonist efficacy (Salehi-Rad *et al*. 2021). In contrast, on administration with quercetin, feeble interaction was observed with 20% decreased expression, suggesting its weak efficacy towards the activation (Salehi-Rad *et al*. 2021).The quantitative data of PPAR-γ expression elicited by QDs of both cell lines are presented in Figure 5 (D).

### PPAR-γ activation inhibits loss of E-cadherin and acquisition of mesenchymal markers-Snail, Slug, Zeb-1 during EMT

EMT is marked as a dreadful reason leading to manifested biological morphogenetic process. Prevailing studies showed that activation PPAR-γ could inhibit EMT markers (Reka *et al*. 2010; Choi *et al*. 2014; Tan *et al*. 2010). Therefore, we sought to determine the effect of PPAR-γ activation on EMT where we treated A549 cells with our synthesized QDs to understand the underlying mechanism in EMT pathway. In the course of EMT process, epithelial markers are seen to be downregulated, whereas mesenchymal markers are seen to be upregulated.^36^ Following the treatment, the expression of E-cadherin was dramatically upregulated by all the QDs (Figure 8 (A)). In comparison to the controls used; the intensity of E-cadherin appeared to be lower than RSG which reflected maximal expression. On analysing the mesenchymal marker expression (Figure 8 (B-F)), the results revealed that the expression of Snail, Slug and Zeb-1 was dramatically downregulated by all the QDs. In contrast, QDs did not have much effect on the expression of N-cadherin and Vimentin. Many studies support our finding wherein quercetin was efficient to inhibit the mesenchymal markers; Snail and Slug, in various human cancers, including lung. The transcription factors are the primary source for regulation of EMT, viz., Snail, Slug, Twist, Zeb-1, N-cadherin, and Vimentin, which behave as a potent repressor for epithelial gene expression (Salehi-Rad *et al*. 2020). In particular, Snail serves as a mediator in promoting the change in adhesion molecule gene expression from E-cadherin to N-cadherin, while Slug act as a substantial target of β-catenin, an important protein possessing a dual role with cell-cell adhesion and gene transcription in epithelial cells. On the other hand, Zeb-1 (zinc finger E-box binding homeobox 1) expresses itself by binding to the E-boxes, followed by repressing some epithelial junctions and activating mesenchymal genes. These zinc-finger proteins have a strong correlation with the transcriptional repression of E-cadherin (Zhuo *et al*. 2008). However, no report illustrates the reason for not affecting N-cadherin and Vimentin. These observations possibly explain that PPAR-γ activation by QDs completely blocks the acquisition of three mesenchymal markers; Snail, Slug, and Zeb-1,while the loss of epithelial marker, E-cadherin was prevented by all the QDs.

**Fig. 8.**
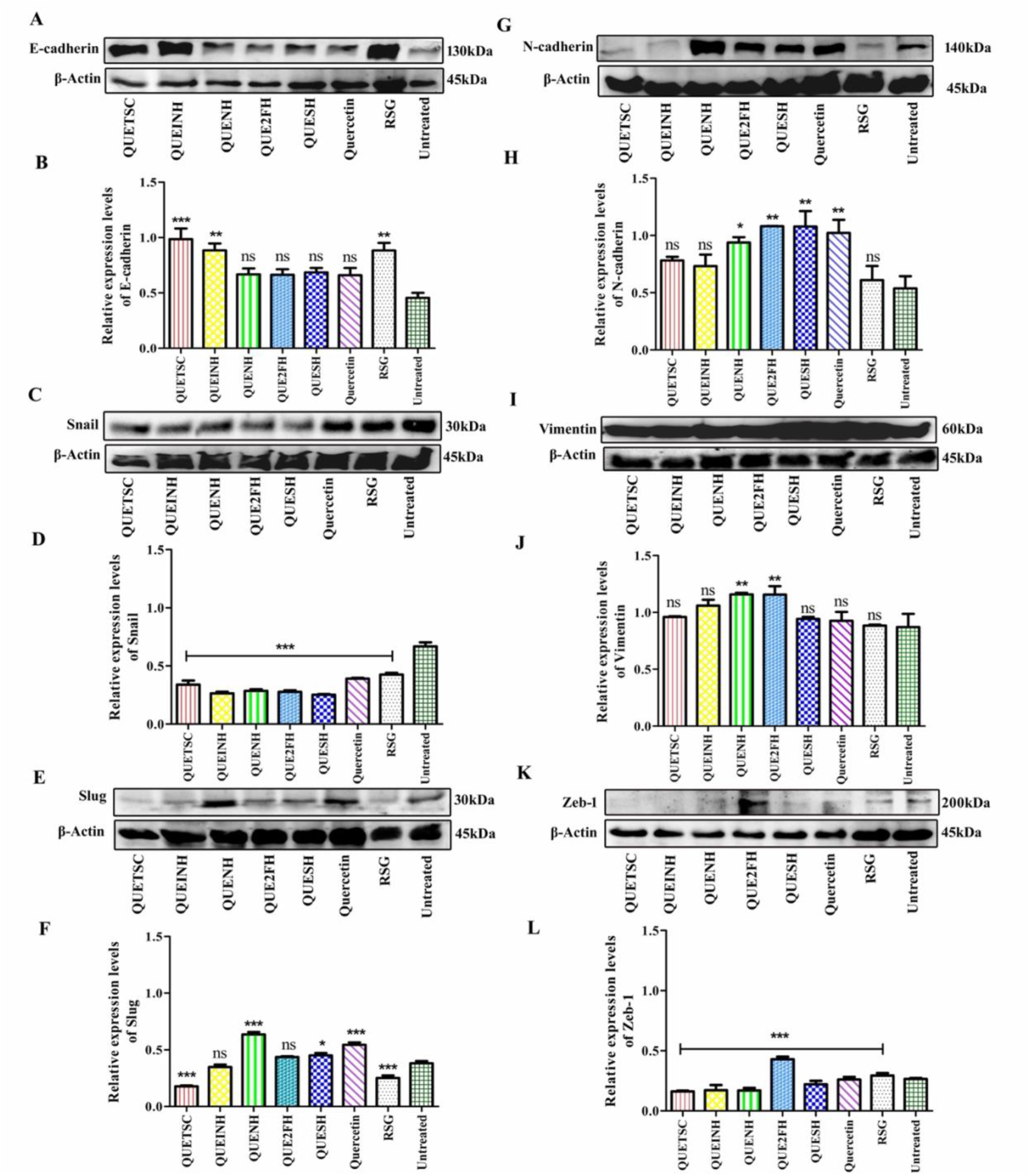
Effect of PPAR-γ ligands (synthesized QDs) on the prevention of epithelial marker and acquisition of mesenchymal markers on A549 cells after 24h of treatment. A549 cells were treated with 1/10^th^ dose of IC_50_ values of QDs, and proteins were obtained, separated on SDS–PAGE, probed with respective antibodies. (B) Representative Western blotting images with their relative protein expression level of (A& B) E-cadherin (C & D) Snail (E & F) Slug (G & H) N-cadherin (I & J) Vimentin (K & L) Zeb-1with respect to β-actin, which serve as equal loading control. Each value is presented as mean±SD for three replicate determinations for each treatment (p<0.05, one-way ANOVA). The error bar represents SD; *p<0.05, **p<0.01 and ***p < 0.001 compared with control group-Untreated cells.

## CONCLUSION

In conclusion, chemical modification of quercetin parent compound led to novel and intrinsically potent partial agonists for PPAR-γ. The present research deduces the central importance of counteracting EMT in A549 cells via PPAR-γ activation through these synthesized QDs. Owing to their drug-like properties; the derivatives exhibited desired effects on inhibiting the growthandEMTinA549 cells at nanomolar concentrations. Of the five derivatives we screened, QUETSC, QUE2FH, and QUESH were found to activate the protein cooperatively with a significantly lower intrinsic activity than that of the full agonist, RSG and the weak agonist, quercetin. Following PPAR-γ partial activation, these three QDs ought to be the most potential to modulate the biochemical markers of EMT with the inhibition of migration invasiveness. These results suggest that QUETSC, QUE2FH, and QUESH appear promising PPAR-γ partial agonists, which can be used as anti-metastatic agent. However, transforming growth factor-beta (TGF-β), a multifunctional cytokine, is a key player to have a greater impact on EMT that deliberately contributes to develop metastasis. Therefore, further investigations are underway to validate the effectiveness of these partial agonists on EMT, induced by TGF-β through partial activation of PPAR-γ.

## Acknowledgment

The authors are thankful to Dr. D. Y. Patil Biotechnology and Bioinformatics Institute, Dr. D. Y. Patil Vidyapeeth, Pune for the physical infrastructure. The authors also acknowledge the Department of Science and Technology Science and Engineering Research Board (DST-SERB), Govt. of India, New Delhi, (File Number: ECR/2016/000943) for financial support and utilizing an optimized supercomputer for dynamics calculations (File Number: YSS/2015/002035). S. Ballav is thankful to DST-SERB for Junior Research Fellowship. Senior Research Fellowship awarded to K. B. Lokhande (Project ID: 2019-3458; File No.: ISRM/11(54)/2019) of the Indian Council of Medical Research (ICMR), New Delhi is also acknowledged. The authors acknowledge the support of the Prof. Subhash Padhye Laboratory, Interdisciplinary Sciences and Technology Research Area Academy (ISTRA), University of Pune, for synthesizing the compounds.

## Declarations

### Ethical Approval

The human NSCLC cell lines A549 and NCI-H460 were procured from the National Centre for Cell Science (NCCS), Pune, Maharashtra, India.

### Consent to Participate

Not applicable

### Consent to Publish

Not applicable

## Authors’ Contributions

SB and AR conceived and designed research. MB and SP were responsible for the synthesis of the compounds. KBL and KVS conducted MD simulation. SaB conducted molecular docking and *in vitro* experiments. SaB analyzed data and wrote the manuscript. SaB, SB, AR and MKP reviewed the manuscript. All authors read and approved the manuscript and all data were generated in-house and that no paper mill was used.

### Funding

This work was supported by Department of Science and Technology Science and Engineering Research Board (DST-SERB),Govt. of India, New Delhi, (File Number: ECR/2016/000943).

### Availability of data and materials

The datasets generated during and/or analysed during the current study are available from the corresponding author on reasonable request.

