## Supplementary figures and tables for "Design, synthesis and biological evaluation of novel quercetin derivatives as PPAR-γ partial agonists by modulating Epithelial-mesenchymal transition in lung cancer metastasis"

### Chromatographic and Spectroscopic Characterization

The structure of quercetin (3,5,7-trihydroxy-2-(3,4-dihydroxyphenyl)-4Hchromen-4-one) comprises of three rings (A,B, and C), 2-phenylchromen-4-one and five hydroxyl groups. Quercetin has its characteristic absorption bands at 3406 and 3283 $cm^{-1}$ for rings A and B respectively (O-H stretching), at 1379 $cm^{-1}$ for phenol (O-H bending), at 1666 $cm^{-1}$ for aryl ketone (C=O stretching), at 1610, 1560, and 1510 $cm^{-1}$ for aromatic ring (C=C stretchings), at 1317 $cm^{-1}$ for in-plane aromatic ring (C-H stretching) and at 933, 820, 679, and 600 $cm^{-1}$ for out-plane aromatic ring (C-H bending) (Catauro *et al.* 2015). In the case of QDs, the strong signal of aryl ketonic C=O confirmed its location between 1655 $cm^{-1}$ to 1754 $cm^{-1}$due to the existence of hydrophilic groups in quercetin at high concentrations. The shoulder bands around 3095 $cm^{-1}$ of QUEINH, QUENH, QUE2FH and QUESH might have contributed to the symmetry and asymmetric stretching of azomethine group (C=N) (Nishat *et al.* 2010) whereas QUETSC exhibited no such bands.

The peaks at 850 $cm^{-1}$ and 570 $cm^{-1}$ for QUEINH, QUENH, QUE2FH and QUESH attributed to the symmetric and asymmetric C-O stretching vibrations and C-CO-C bending (Figure S2). Whilst a shift from lower wavenumber region was witnessed for C-O band and C-CO-C bending in QUETSC at 878 $cm^{-1}$ and 593 $cm^{-1}$, designating the absence of aromatic ring in the core part of thiosemicarbazine, thus alternating the microenvironment of quercetin. These findings account the fact that the upshift in terms of energy absorption of the O-H bending band and the ketone band is a characteristic of hydrogen bonded O-H groups (Catauro *et al.* 2015). The spectrum of all the QDs shows a strong band in the region of 873 to 934 $cm^{-1}$due to N-N mode that leads to reduced repulsion between the adjacent nitrogen atoms that consists of lone pairs (Dash *et al.* 2008).Abrupt changes at 1587 $cm^{-1}$ and 1497 $cm^{-1}$ were noticed in the absorption network of the double bond for QUEINH, which may be probably due to the heterocyclic interaction of ions of isonicotinic acid hydrazine with quercetin. Furthermore, in all the spectras of QDs, the peak across 1354 $cm^{-1}$ wavenumber was due to O-H bending vibrations in the hydration water. The broad band of O-H stretching demonstrates the strength of hydrogen bond within quercetin compound resulting from the strong overlapping of hydroxyl group of quercetin with the -NH group of all the QDs. These outcomes club the strength of interionic hydrogen bonding of QDs, unlike the structure of QUETSC which lacks an aromatic ring in its derivative, thus leading to a weak interaction with quercetin.

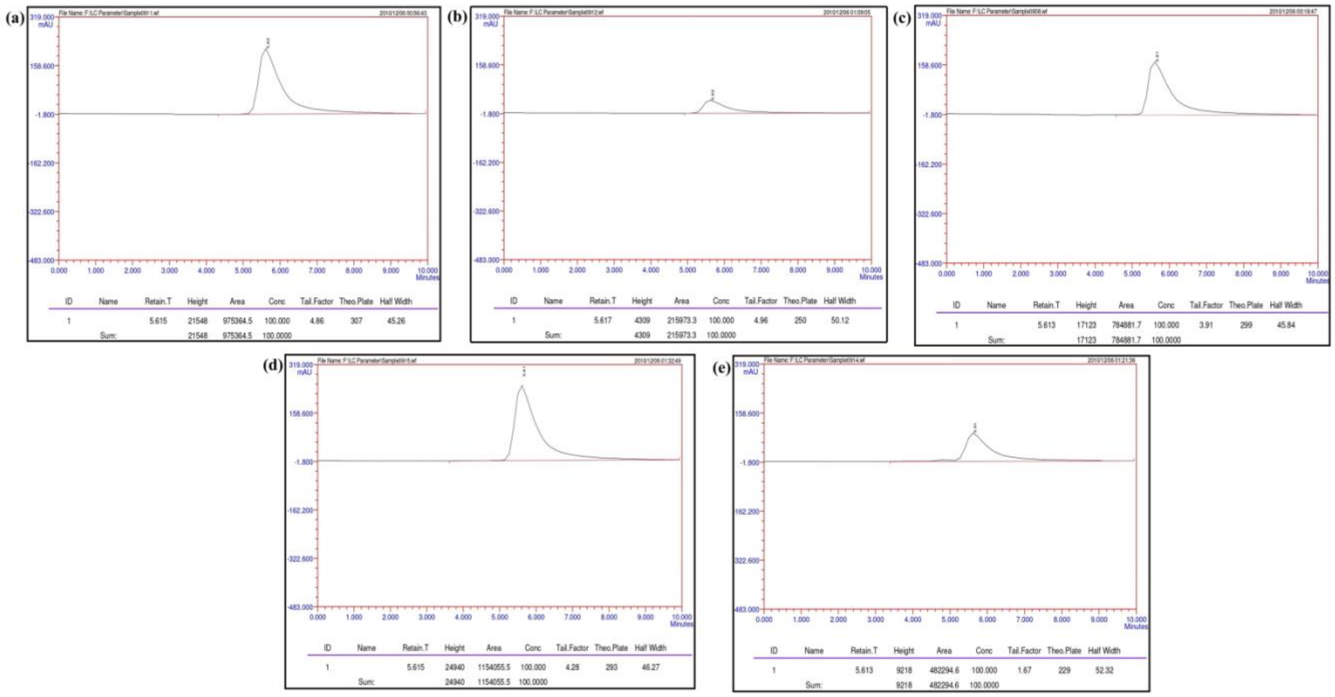


**Fig. S1 HPLC Chromatogram of newly synthesized QDs. The fraction at various retention time ($\mathbf{t_R}$) corresponds to** (a)QUETSC-$\mathbf{t_R}$ = 5.62; (b) QUEINH-$\mathbf{t_R}$ = 5.62; (c) QUENH-$\mathbf{t_R}$ = 5.61; (d)QUE2FH - $\mathbf{t_R}$ = 5.62 ; (e) QUESH -$\mathbf{t_R}$ = 5.62**.** The peaks of QDs were well separated at different retention times. No interferences or excipient peaks co eluted with the analytes, indicating the method is selective and specific in relation to the medium and excipients used in this study.

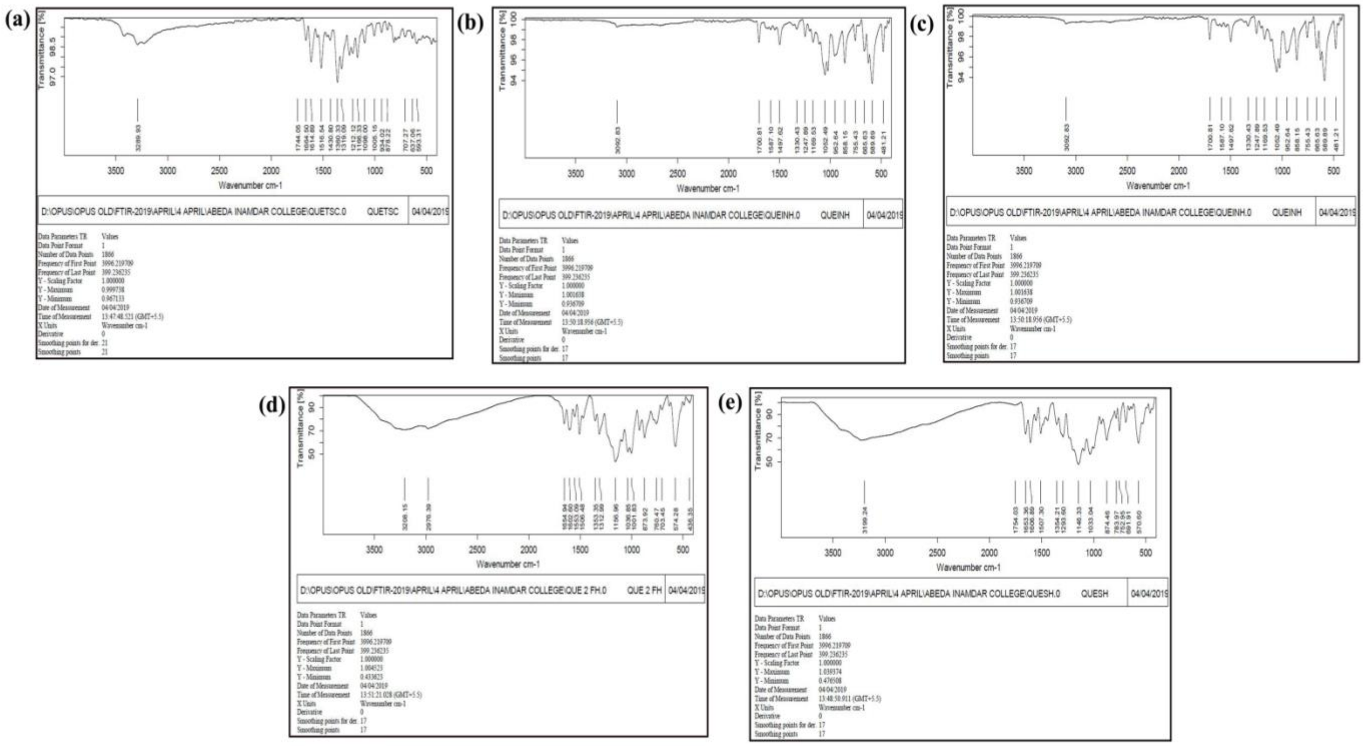


**Fig. S2 FT-IR Spectra depicting the presence of functional groups in newly synthesized QDs.** (a) QUETSC; (b) QUEINH; (c)QUENH; (d)QUE2FH; (e)QUESH

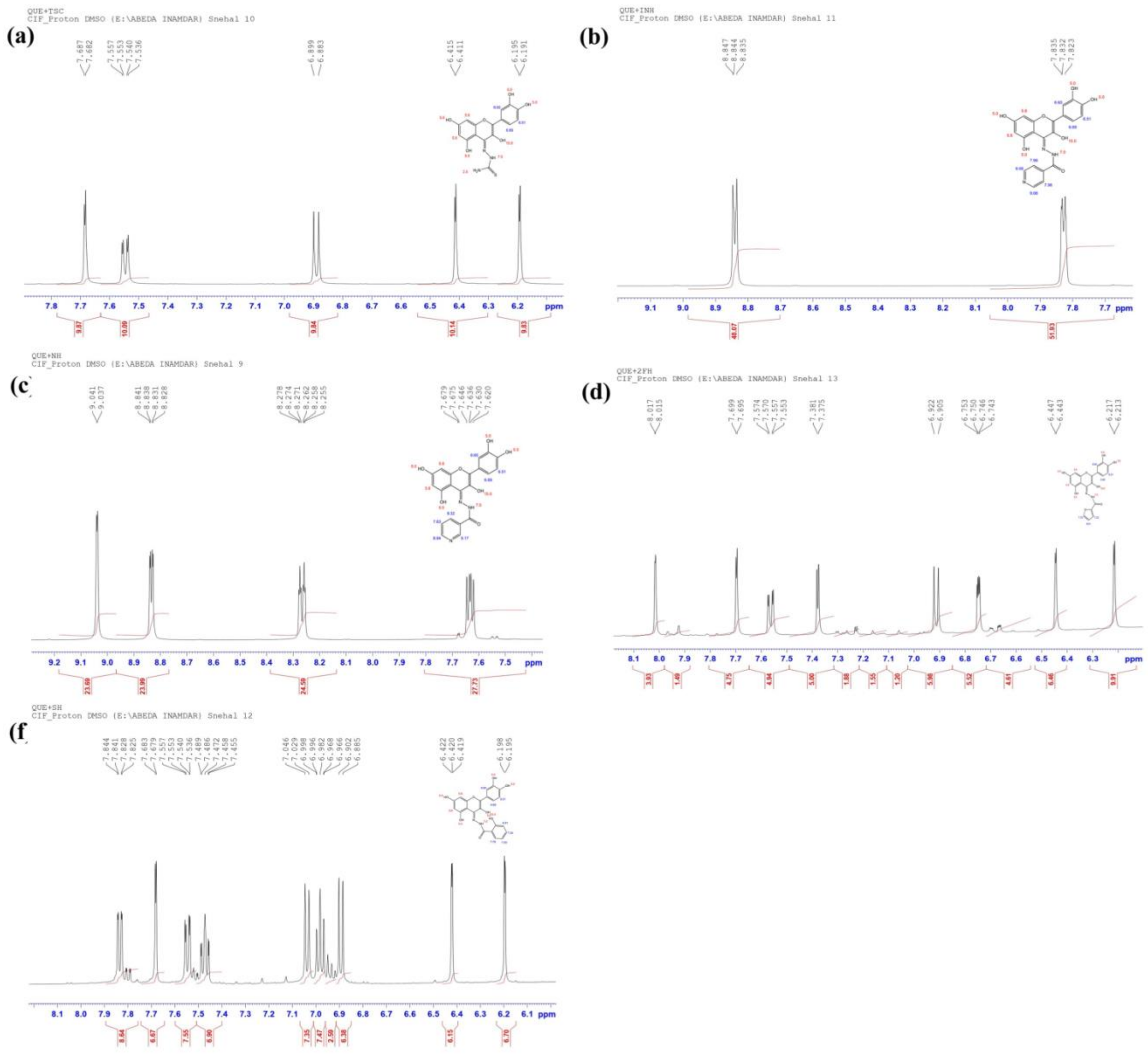


**Fig. S3 NMR H1 Spectra determining the purity and molecular structure of the synthesized QDs.** (a)QUETSC; (b) QUEINH; (c)QUENH; (d)QUE2FH; (e)QUESH

### ADME Prediction profiles of QDs

Table S1 summarizes the detailed values of each parameter of Lipinski's rule retrieved for all the five QDs. Our analysis showed that all the tested entities passed the Lipinski screening test albeit for one factor; the HBD which exceeded the limit above 5.All the five QDs were noted to be within the acceptable range for ADME pharmacokinetic parameters. Another pharmacokinetic property was observed that they all are not P-gp substrate, which makes them a successful drug against multidrug-resistant cancer cells, overexpressing this drug transporter. The prediction of putative drug–drug interaction through Cytochromes P450 (CYPs) inhibition of all the compounds have also been reported in Table S2.We sighted that almost all our compounds fall in the optimal range for each property.

**Table S1 The molecular descriptors and Lipinski properties of synthesized QDs ligands using SwissADME web tool**

| Name of the compound | Log P (<5) | MW in g/mol (<500Da) | HBA(<10) | HBD(<5) | Molar refractivity (<130) |
|---|---|---|---|---|---|
| QUETSC | 1.23 | 375.36 | 7 | 7 | 96.49 |
| QUEINH | 1.21 | 421.38 | 9 | 6 | 108.33 |
| QUENH | 0.89 | 421.36 | 9 | 6 | 108.7 |
| QUE2FH | 1.56 | 410.33 | 9 | 6 | 103.17 |
| QUESH | 1.75 | 436.37 | 9 | 7 | 112.93 |

**Table S2 Physicochemical, pharmacokinetics properties and drug-likeness of synthesized QDs using to SwissADME web tool.**

| Name of the compound | Pgp substrate | CYP1A2 inhibitor | CYP2C19 inhibitor | CYP2C9 inhibitor | CYP2D6 inhibitor | CYP3A4 inhibitor |
|---|---|---|---|---|---|---|
| QUETSC | No | Yes | No | No | No | No |
| QUEINH | No | No | No | No | No | No |
| QUENH | No | Yes | No | No | No | No |
| QUE2FH | No | Yes | No | Yes | No | No |
| QUESH | No | Yes | No | No | No | No |

**Cell cycle analysis**

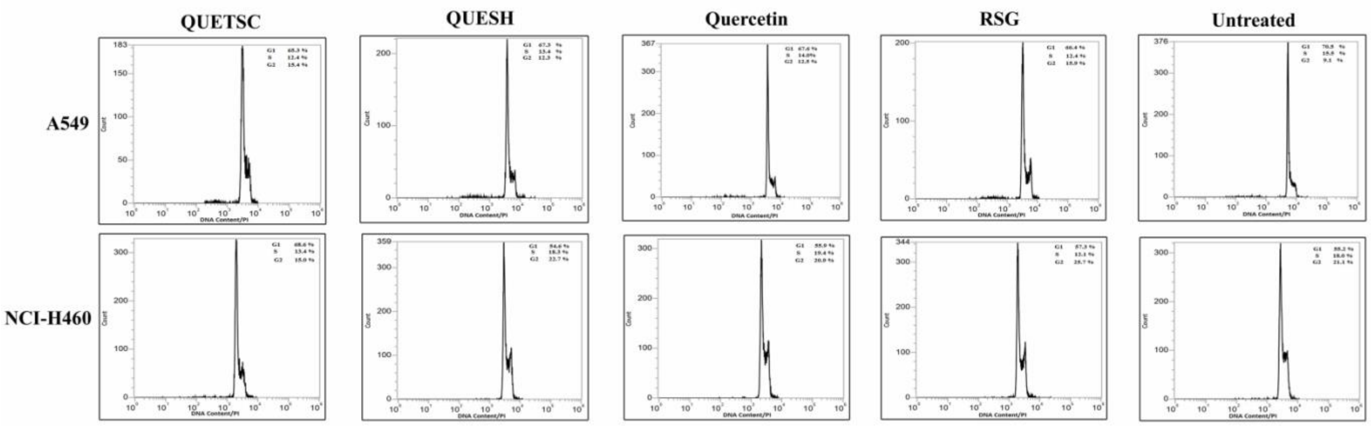


**Fig. S4 Cell cycle analysis at 48h of A549 and NCI-H460 cells.** Cells were incubated with $IC_5$ doses of QUETSC, QUESH, quercetin and RSG for 48 h, and then collected, fixed, stained with propidium iodide and analyzed for DNA content by flow cytometry. Percentages of cells in different phases of cell cycle: G1, G2 and S are indicated in each panel.

**Scratch assay**

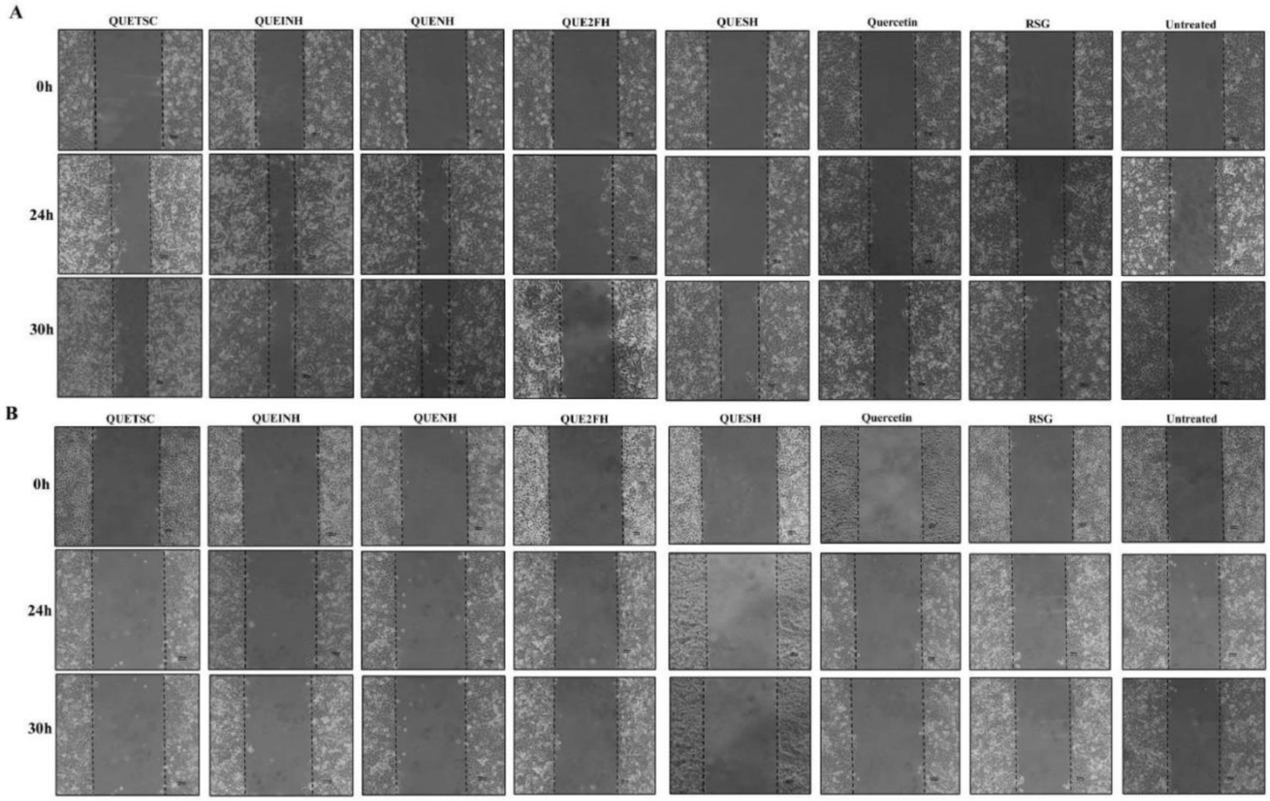

**Fig. S5 PPAR-γ Based Ligands- Synthesized QDs inhibits migration of A549 and not in NCI-H460 cells.** A549 (A) and NCI-460 (B) cells were seeded in 35 mm plate and cultured for 24h in normal medium followed by serum starvation of 24 h before scratching. With the help of sterile 2 µl micropipette tip, a single scratch was made in each plate and subsequently treated with $IC_5$ doses of QDs for 30h under standard conditions.
