## Supplementary figures and images for "Design, synthesis and biological evaluation of novel quercetin derivatives as PPAR-γ partial agonists by modulating Epithelial-mesenchymal transition in lung cancer metastasis"

### Supplementary Figure 1

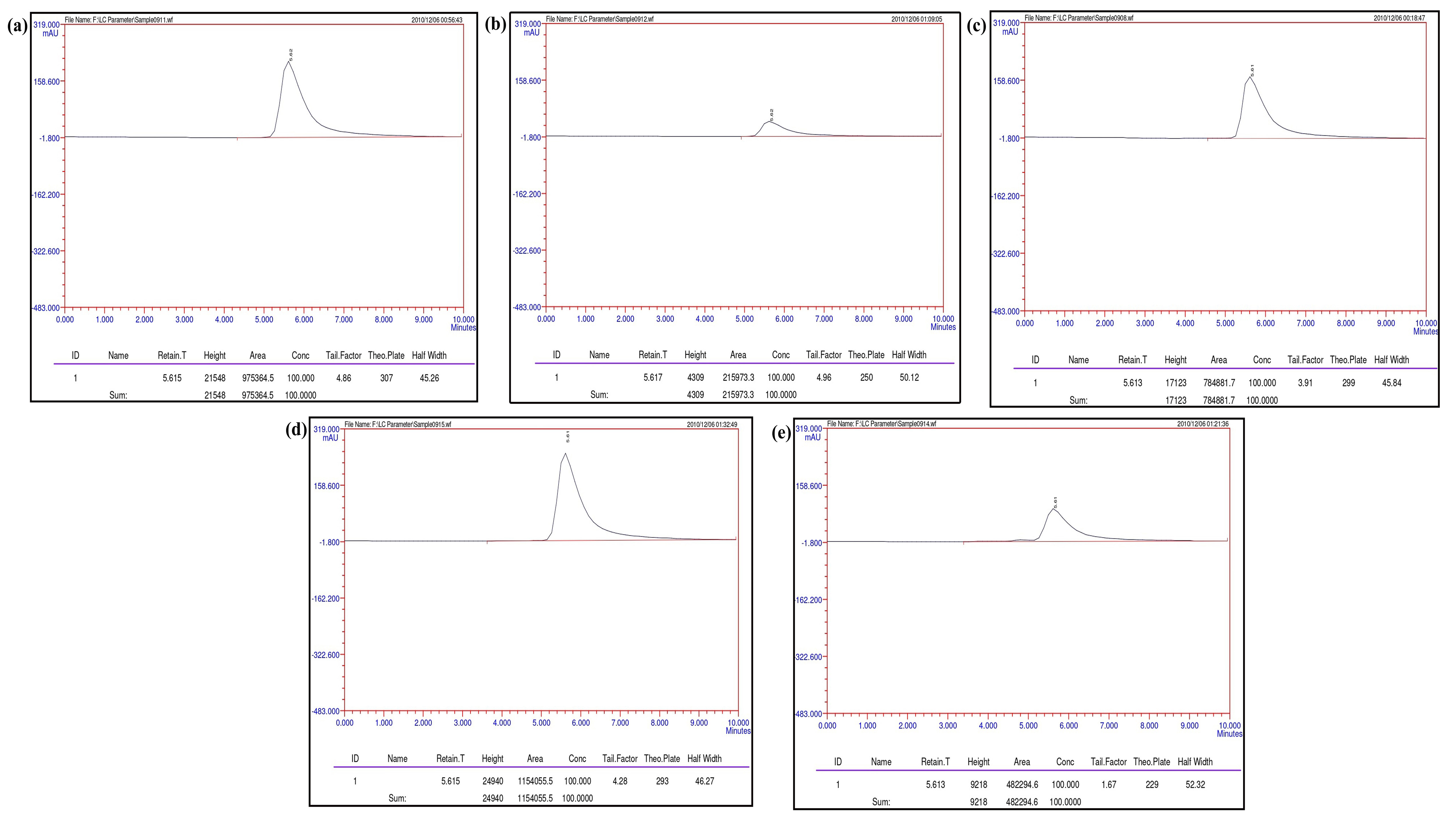

### Supplementary Figure 2

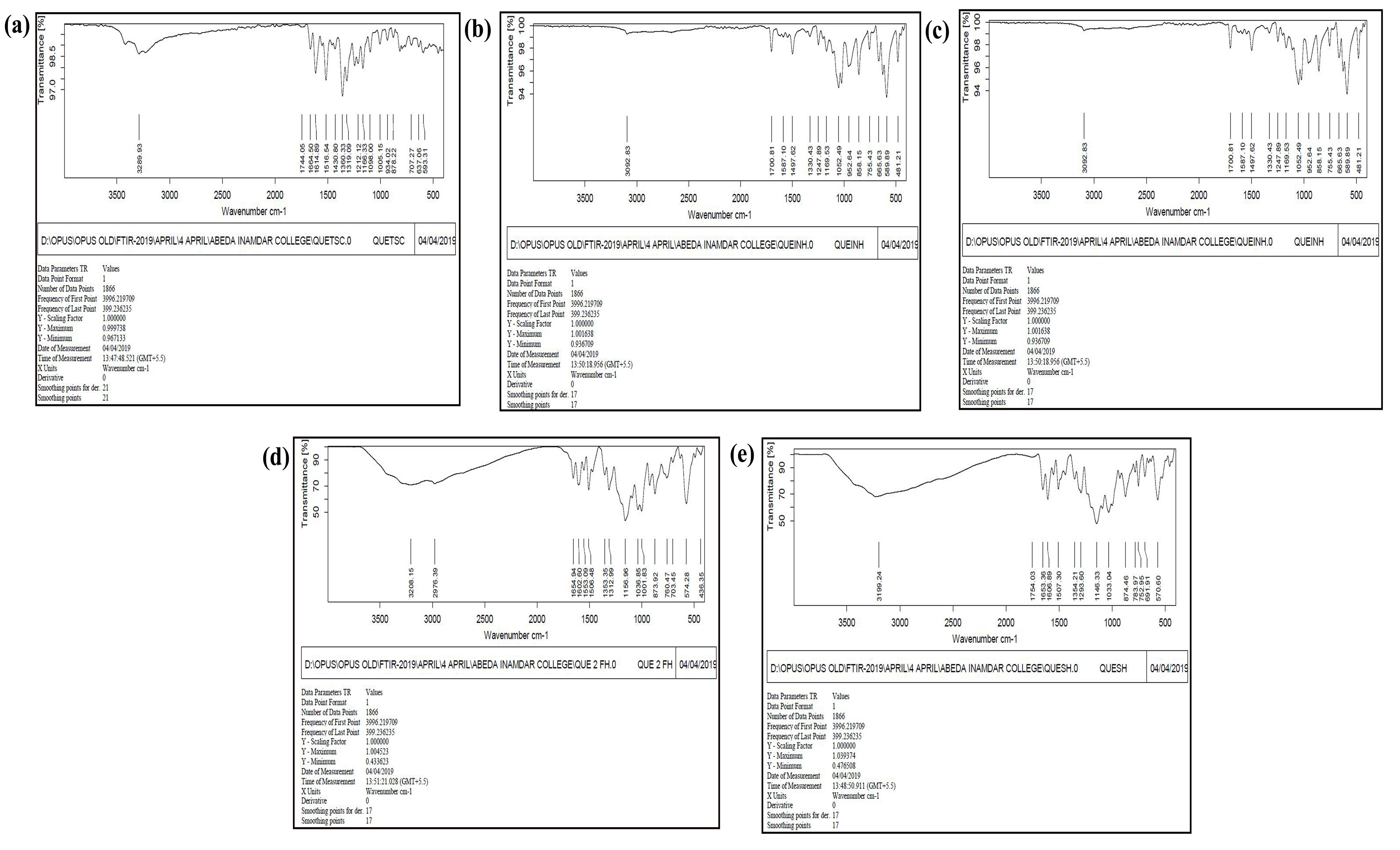

### Supplementary Figure 3

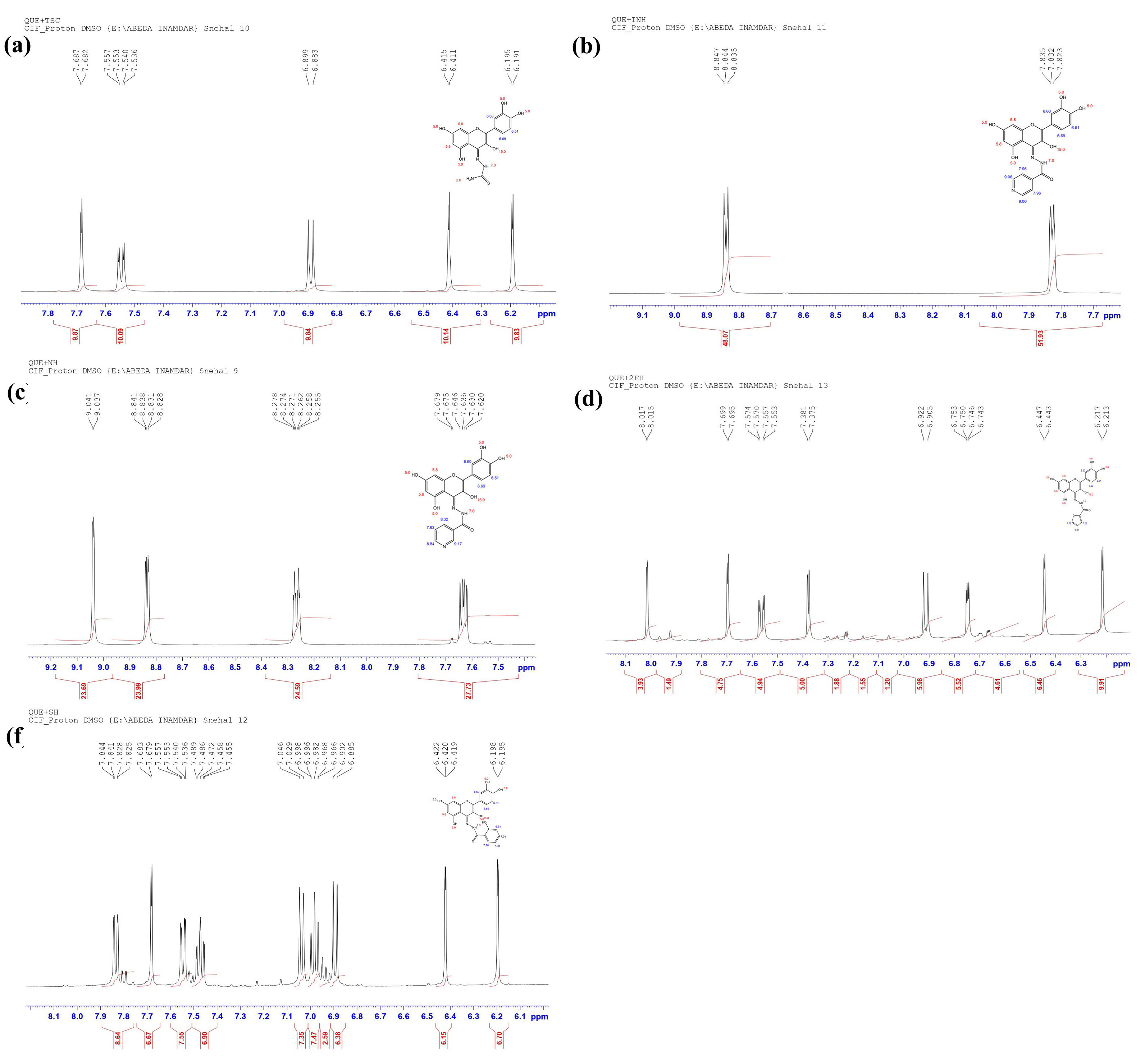

### Supplementary Figure 4

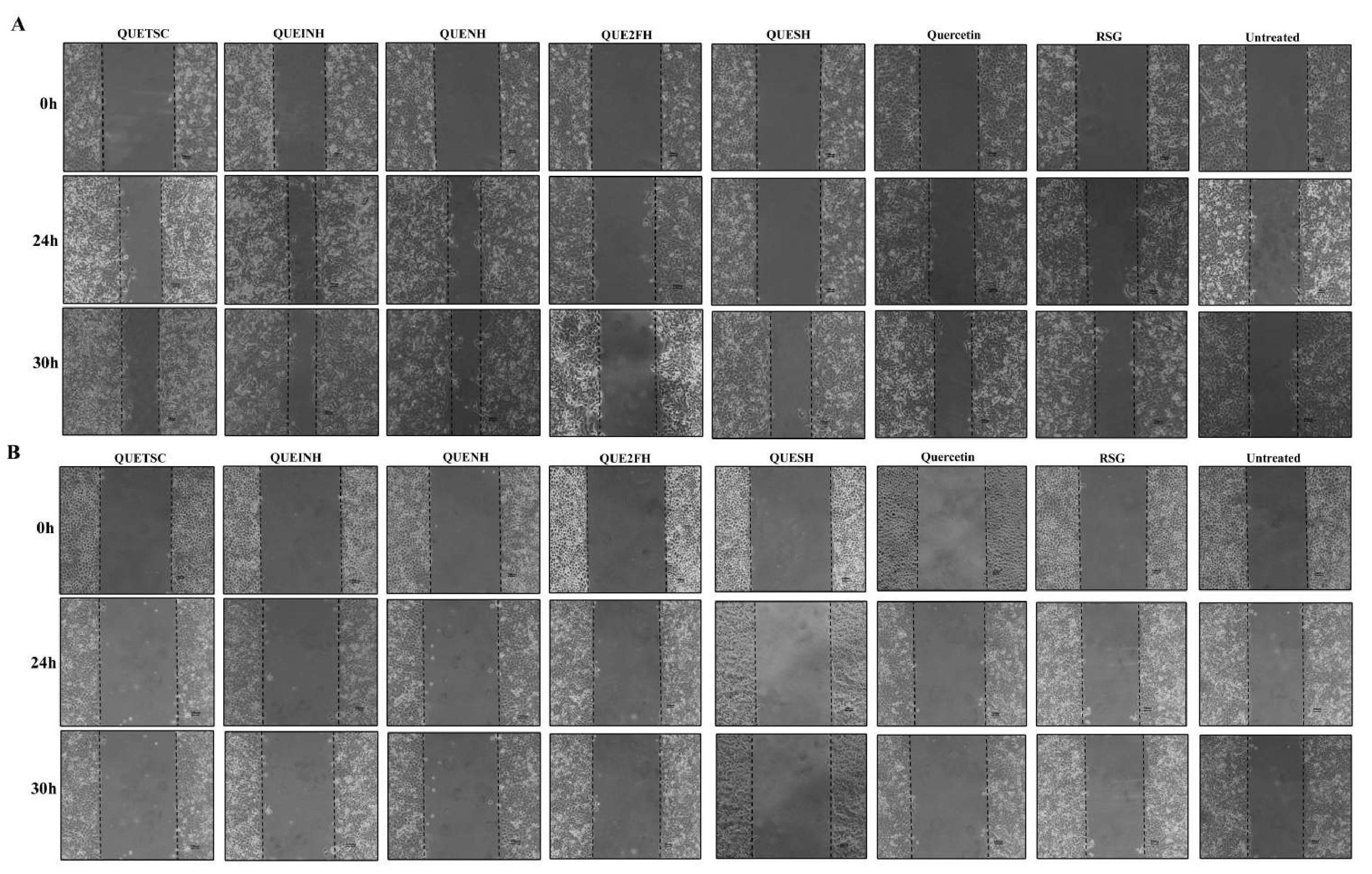

### Supplementary Figure 5

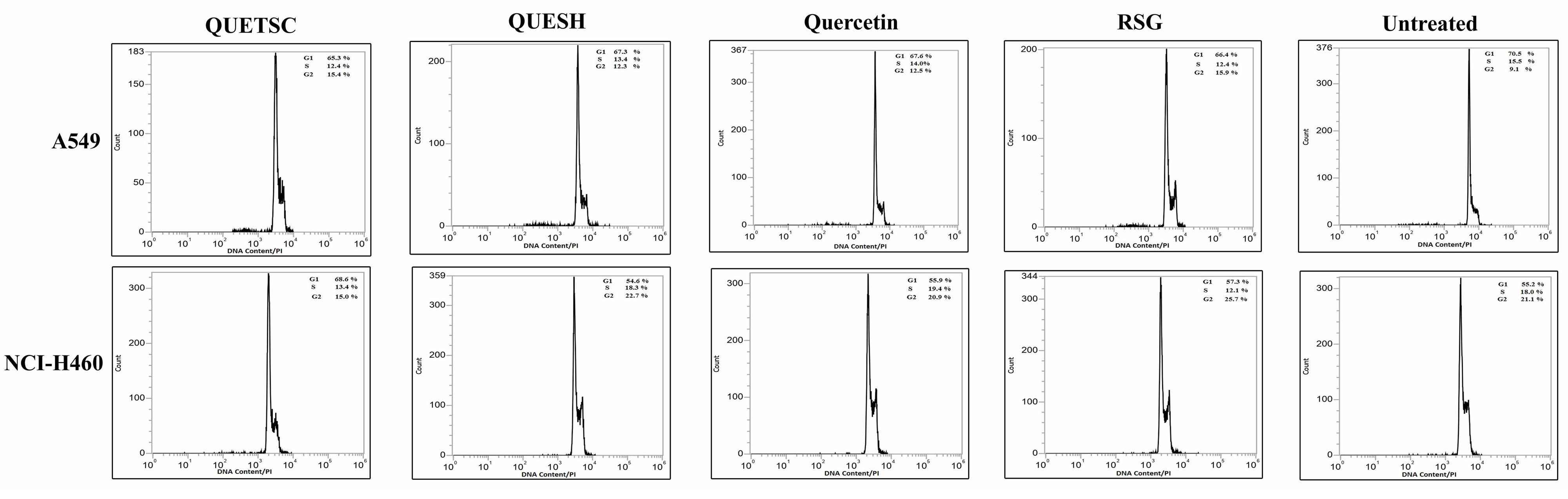
